# Characteristic and quantifiable COVID-19-like abnormalities in CT- and PET/CT-imaged lungs of SARS-CoV-2-infected crab-eating macaques (*Macaca fascicularis*)

**DOI:** 10.1101/2020.05.14.096727

**Authors:** Courtney L. Finch, Ian Crozier, Ji Hyun Lee, Russ Byrum, Timothy K. Cooper, Janie Liang, Kaleb Sharer, Jeffrey Solomon, Philip J. Sayre, Gregory Kocher, Christopher Bartos, Nina M. Aiosa, Marcelo Castro, Peter A. Larson, Ricky Adams, Brett Beitzel, Nicholas Di Paola, Jeffrey R. Kugelman, Jonathan R. Kurtz, Tracey Burdette, Martha C. Nason, Irwin M. Feuerstein, Gustavo Palacios, Marisa C. St. Claire, Matthew G. Lackemeyer, Reed F. Johnson, Katarina M. Braun, Mitchell D. Ramuta, Jiro Wada, Connie S. Schmaljohn, Thomas C. Friedrich, David H. O’Connor, Jens H. Kuhn

**Affiliations:** Integrated Research Facility at Fort Detrick, National Institute of Allergy and Infectious Diseases, National Institutes of Health, Fort Detrick, Frederick, MD 21702, USA; Integrated Research Facility at Fort Detrick, Clinical Monitoring Research Program Directorate, Frederick National Laboratory for Cancer Research supported by the National Cancer Institute, Frederick, MD 21702, USA; Center for Infectious Disease Imaging, Warren G Magnuson Clinical Center, National Institutes of Health, Bethesda, MD, 20814, USA; United States Army Medical Research Institute of Infectious Diseases, Fort Detrick, Frederick, Maryland 21702, USA; Biostatistics Research Branch, National Institute of Allergy and Infectious Diseases, National Institutes of Health, Rockville, MD 20892, USA; Department of Pathobiological Sciences, University of Wisconsin-Madison, Madison, WI 53706, USA; Department of Pathology and Laboratory Medicine, University of Wisconsin-Madison, Madison, WI 53706, USA; Wisconsin National Primate Research Center, Madison, WI 53706, USA

## Abstract

Severe acute respiratory syndrome coronavirus 2 (SARS-CoV-2) is causing an exponentially increasing number of coronavirus disease 19 (COVID-19) cases globally. Prioritization of medical countermeasures for evaluation in randomized clinical trials is critically hindered by the lack of COVID-19 animal models that enable accurate, quantifiable, and reproducible measurement of COVID-19 pulmonary disease free from observer bias. We first used serial computed tomography (CT) to demonstrate that bilateral intrabronchial instillation of SARS-CoV-2 into crab-eating macaques (*Macaca fascicularis*) results in mild-to-moderate lung abnormalities qualitatively characteristic of subclinical or mild-to-moderate COVID-19 (e.g., ground-glass opacities with or without reticulation, paving, or alveolar consolidation, peri-bronchial thickening, linear opacities) at typical locations (peripheral>central, posterior and dependent, bilateral, multi-lobar). We then used positron emission tomography (PET) analysis to demonstrate increased FDG uptake in the CT-defined lung abnormalities and regional lymph nodes. PET/CT imaging findings appeared in all macaques as early as 2 days post-exposure, variably progressed, and subsequently resolved by 6–12 days post-exposure. Finally, we applied operator-independent, semi-automatic quantification of the volume and radiodensity of CT abnormalities as a possible primary endpoint for immediate and objective efficacy testing of candidate medical countermeasures.

---

The causative agent of human coronavirus disease 2019 (COVID-19), severe acute respiratory syndrome coronavirus 2 (SARS-CoV-2), likely emerged in Wǔhàn, Húběi Province, China in November 2019 (1–3). The virus rapidly spread through the human population, causing more than 4.2 million infections and almost 300,000 deaths globally by May 14, 2020 (4). Infections result in a wide spectrum of disease ranging from asymptomatic to mild upper respiratory illness to severe pneumonia that can progress to acute respiratory distress syndrome (ARDS) and death despite aggressive supportive care (5). After a short but variable incubation period, most patients with COVID-19 develop self-limiting fever, cough, nonspecific fatigue, and myalgia (6–11). Some patients develop non-productive cough and dyspnea related to lower respiratory tract involvement; particularly in patients of older age or with co-morbidities, this involvement can lead to severe, progressive disease and unfavorable outcomes (5). Well-documented characteristic lung CT findings in humans include ground-glass opacities (GGOs) with or without reticulation, reticulonodular opacities, inter- or intralobular septal paving, or consolidation in a bilateral, lobar to sub-segmental, and peripheral distribution (*6–8*, 12). Notably, GGOs have been described in patients who are shedding SARS-CoV-2 but do not present with clinical signs (13, 14). Bilateral diffuse alveolar damage, type II pneumocyte hyperplasia, interstitial fibrosis and inflammation, hemorrhage, and edema with syncytia appear to be typical lung histopathological findings seen in a limited human data set that also suggests a high rate of venous thromboembolism (15–19).

Currently available rodent/carnivore/tree shrew (20–24) and nonhuman primate (NHP) (*11*, 25–29) models of SARS-CoV-2 infection do not accurately reflect severe human COVID-19. NHPs, considered an evolutionary proximate for human disease modeling, develop no or only mild clinical disease signs (11, 26–29). In SARS-CoV-2-infected rhesus monkeys (*Macaca mulatta*), quantifiable virus shedding, virus-specific immune responses, and limited histopathologic lesions have been observed (*11, 25*, 27–29). However, in both human disease and animal models, the temporal and mechanistic relationship between viral replication, subsequent immunopathology (30, 31), and clinical disease remains uncertain. Furthermore, in the available NHP models, all of which are sublethal, markers of clinical disease (cage-side scoring, chest X-ray) have been of limited sensitivity. More concerningly, both metrics are subject to observer bias (32–35).

Reliable animal models needed for rapid development and evaluation of candidate medical countermeasures (MCMs) require an unbiased reproducible and quantifiable metric of disease that mirrors key aspects of COVID-19. Based on the rather limited X-ray findings in the lungs of reported NHP models of SARS-CoV-2 infection with either mild or no clinical signs (11, 25, 27–29), we turned to high-resolution chest CT and PET/CT to characterize lung abnormalities in infected NHPs toward longitudinal quantitative comparison.

We used direct bilateral primary intrabronchial instillation in a 1-day-apart staggered design to expose two groups of three crab-eating macaques (*Macaca fascicularis*) to medium (mock group macaques M1–3) or medium including 3.65×10^6^ pfu/macaque of SARS-CoV-2 (virus group macaques V1–3) (**Supplementary Table 1**). All macaques were observed daily for 11 days prior to exposure (day [D] 0) and for 30 days post-exposure. Physical examination scores and blood, conjunctival, nasopharyngeal, oropharyngeal, rectal, fecal, and urine specimens were collected at identical timepoints. Virus-exposed macaques were indistinguishable from mock group macaques during the pre-exposure time period. Two pre-exposure chest CT and whole-body 2-deoxy-2-[^18^F]-fluoro-D-glucose (FDG) PET/CTs and eight post-exposure CTs and three post-exposure PETs were performed at identical timepoints (**Figure 1**). In line with previously published results (11, 25–29), none of the macaques developed any major clinical abnormalities (including by cage-side assessment and clinical scoring or physical examination) throughout the study and clinical laboratory results were not significantly different between the mock-exposed and virus-exposed groups (**Supplementary Tables 2–3**). SARS-CoV-2 RNA could not be detected by RT-qPCR in any sample from mock-exposed macaques but was variably present during the early days post-exposure in conjunctival, fecal, nasopharyngeal, oral, and rectal swabs, but never in plasma or urine (**Figure 2a**). Anti-SARS-CoV-2 IgG antibodies were not detectable by ELISA in mock-exposed macaques but were detectable at D10 post-exposure and continued to rise in all virus-exposed macaques to at least D19 (**Figure 2b**). Consistent with ELISA results, fluorescent neutralization titers generated from sera were undetected until D10 and were detected only in virus-exposed macaques (**Figure 2c**). Longitudinal measurement of selected peripheral cytokines revealed between- and within-group differences with marked abnormalities noted in macaque V3, which also had the highest IgG antibody titers **(Supplementary Figure 1)**.

**Figure 1.**
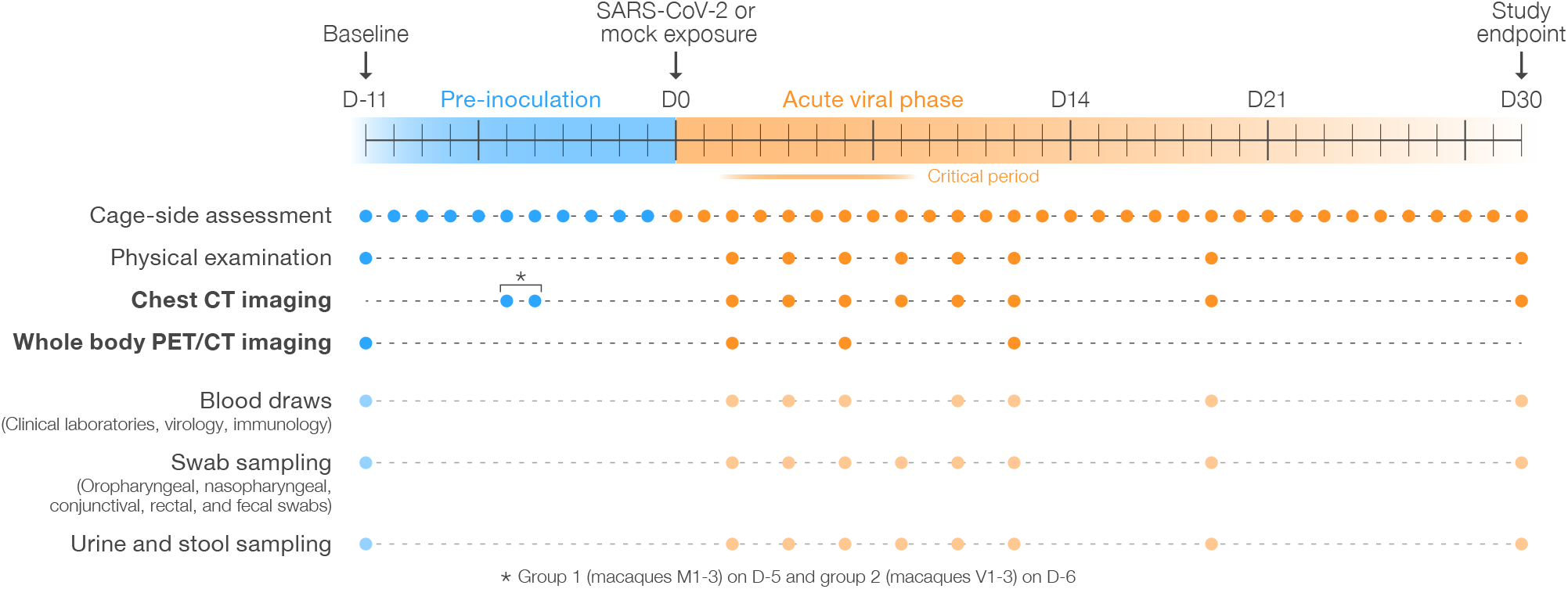
Study overview. Three crab-eating macaques (*Macaca fascicularis*) were exposed to ≈3.65×10^6^ pfu SARS-CoV-2 each by direct bilateral primary post-carinal intrabronchial instillation and sampled as outlined. A second group of three crab-eating macaques was mock-exposed and sampled in the same manner one day prior to the virus-exposed group (exception: a second baseline CT image was recorded for each group on a single day, D-6 or D-5 (*)).

**Figure 2.**
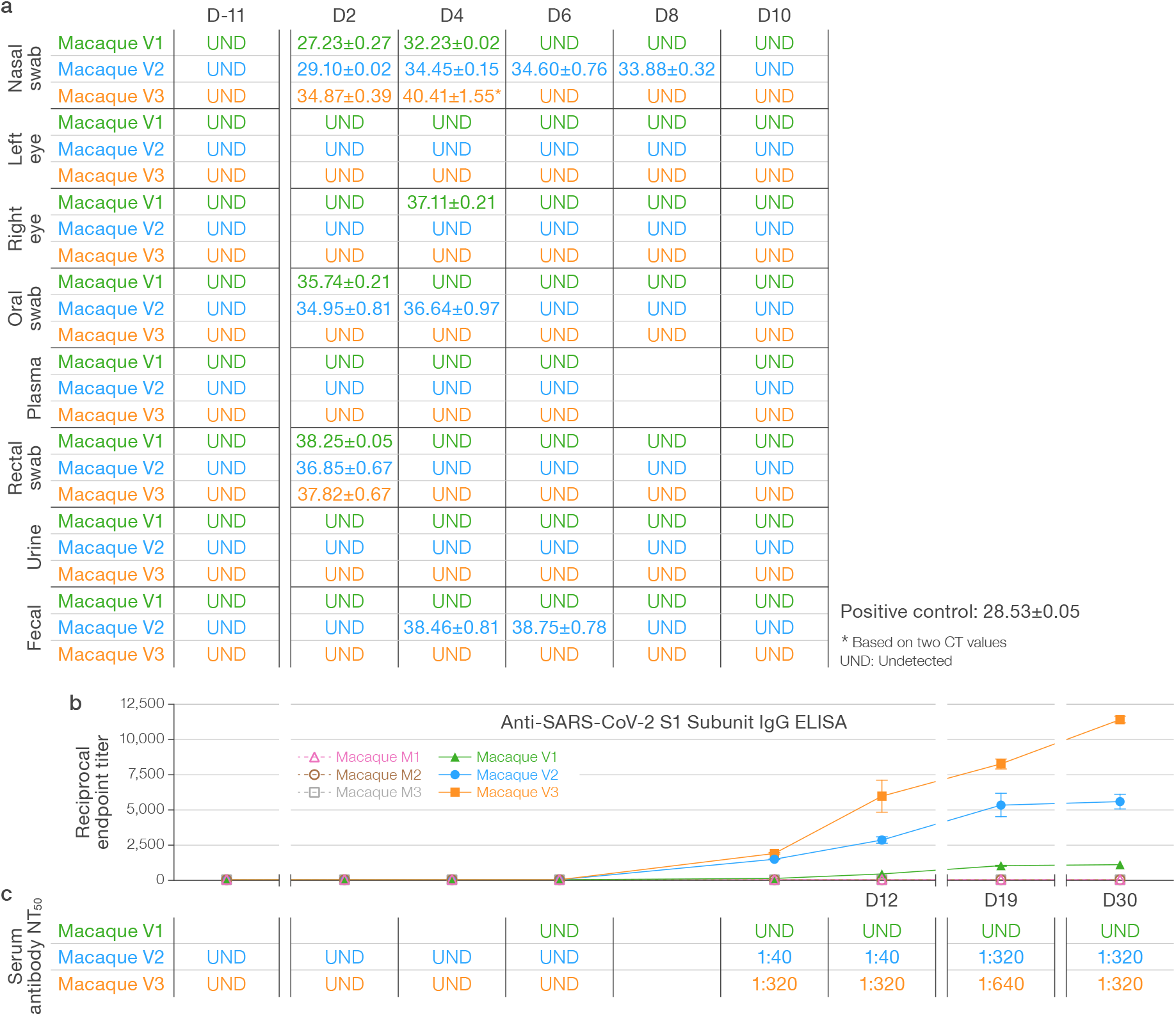
Detection of SARS-CoV-2 RNA and specific immune responses in SARS-CoV-2-inoculated and mock-inoculated macaques. **a)** RT-qPCR targeting the SARS-CoV-2 N protein was performed from baseline through D10 in nasopharyngeal swabs for macaques of both groups. **b)** Anti-SARS-CoV-2 S1 subunit IgG ELISA results are expressed as reciprocal endpoint titers over time for both mock group (M) and virus group (V) macaques. **c)** Fluorescence neutralization assays were performed on sera from all macaques on all days. NT_50_, half-maximal neutralization titer.

With the exception of minor and transient abnormalities on baseline imaging, CT scans of all mock-exposed macaques appeared generally normal over the entire study period (**Supplementary Figure 2)**. However, all virus group macaques developed lung abnormalities clearly visible by chest CT as early as D2. Qualitatively, the distribution morphology, and duration of abnormalities described a spectrum similar to mild-moderately ill humans with COVID-19. Characteristic CT findings in all virus group macaques included bilateral peripheral GGOs variably in association with intra- or interlobular septal prominence (so-called “crazy paving”), reticular or reticulonodular opacities, peri-bronchial thickening, subpleural nodules, and, in one macaque, dense alveolar consolidation with air bronchograms (**Figures 3–4a, Videos 1–3, Supplementary Figure 3).** Longitudinal serial CT scans showed heterogeneity in the duration and evolution of these abnormalities over the next week from rapid improvement within a few days (macaque V1) to persistence and progression (macaques V2, V3) **(Figure 4a, Supplementary Figure 3).** Nonetheless, by D19, chest CT universally showed complete or nearly complete resolution of lung abnormalities in all virus group macaques. Individual and per-group average radiologist-derived CT scores (adapted from a scoring system generated from human COVID-19 CT images) demonstrate the extent and duration of these qualitative findings **(Figure 5).**

**Figure 3.**
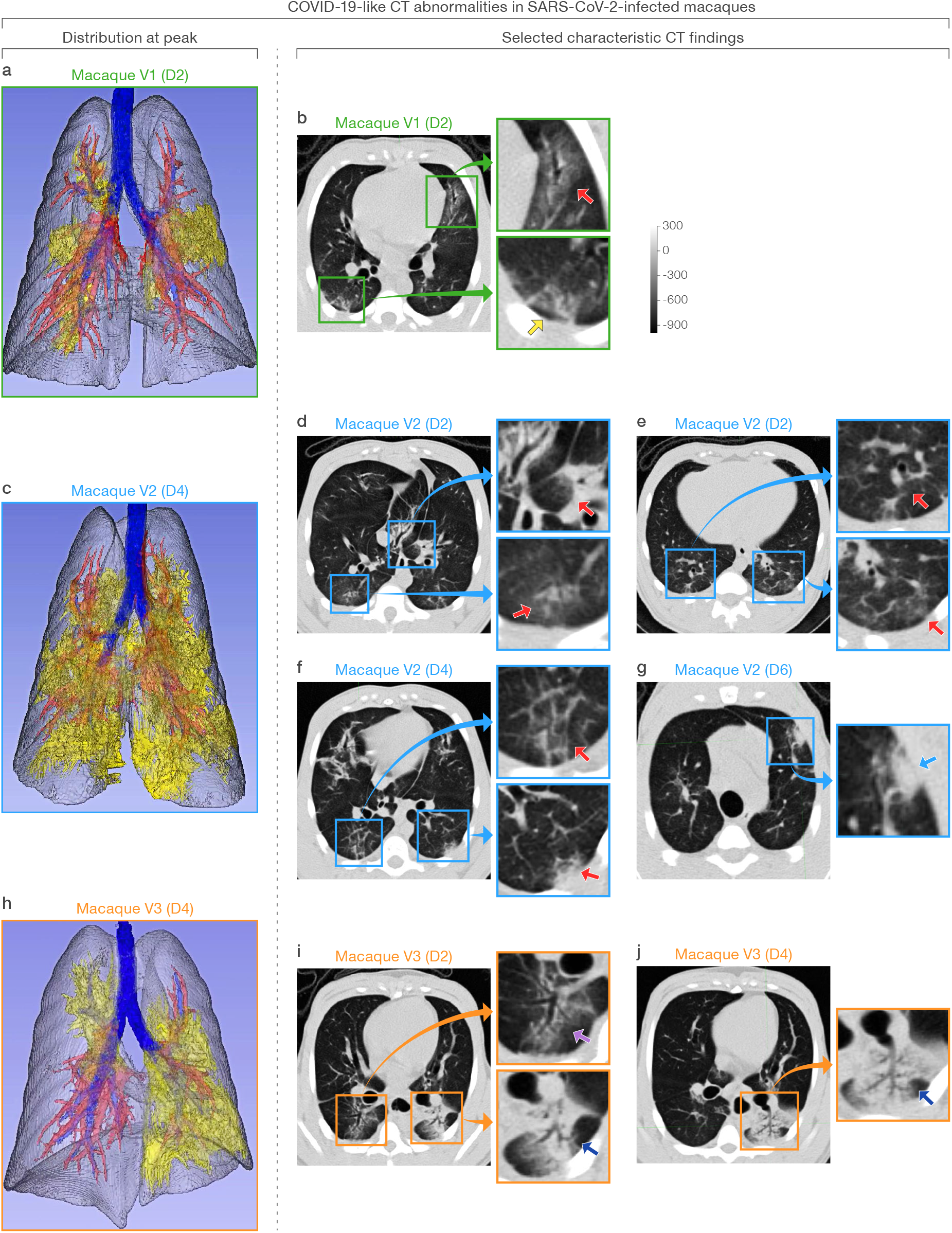
COVID-19-like CT abnormalities in the lungs of SARS-CoV-2 inoculated macaques V1 (a–b), V2 (c–g), and V3 (h–j). Distribution of CT scan abnormalities in 3D images of SARS-CoV-2-inoculated macaque V1 **(a)**, V2 **(c)**, and V3 **(h)**. Blue: airways; gray: normal lung; red: vessels; yellow: imaging abnormalities. **b)** Selected characteristic abnormalities in macaque V1 include peripheral, peri-bronchial ground-glass opacity (GGO) in the left middle lobe (green inset, top, red arrow) and peripheral GGO with reticulation in the posterior right lower lobe (green inset, bottom, yellow arrow). **d–g)** Selected characteristic abnormalities in macaque V2 include **d)** peri-bronchial consolidation in the left accessory lobe (blue inset, top, red arrow) and posterior GGO with reticulation in the posterior right lung (blue inset, bottom, red arrow), **e)** bilateral posterior GGO with reticulation (blue inset, top and bottom, red arrows), **f)** GGO with superimposed paving (interlobular septal thickening) in right posterior lung (blue inset, top) and mixed GGO with pleural-based consolidation in left posterior lung (blue inset, bottom), and **g)** pleural-based mixed GGO and consolidation developing on D6 (blue inset, blue arrow). **i–j)** Selected characteristic abnormalities in macaque V3 include **i)** GGO with air bronchogram in right posterior lung (orange inset, top, purple arrow), alveolar consolidation with peripheral GGO and air bronchogram in left posterior lung (orange inset, bottom, blue arrow), and **j)** expanding and more dense alveolar consolidation with air bronchogram on D4 (orange inset, blue arrow). Representative 3D-rendered videos of **a)**, **c)**, and **h)** demonstrating whole-lung pathology are shown in **Videos 1–3**, respectively.

**Figure 4.**
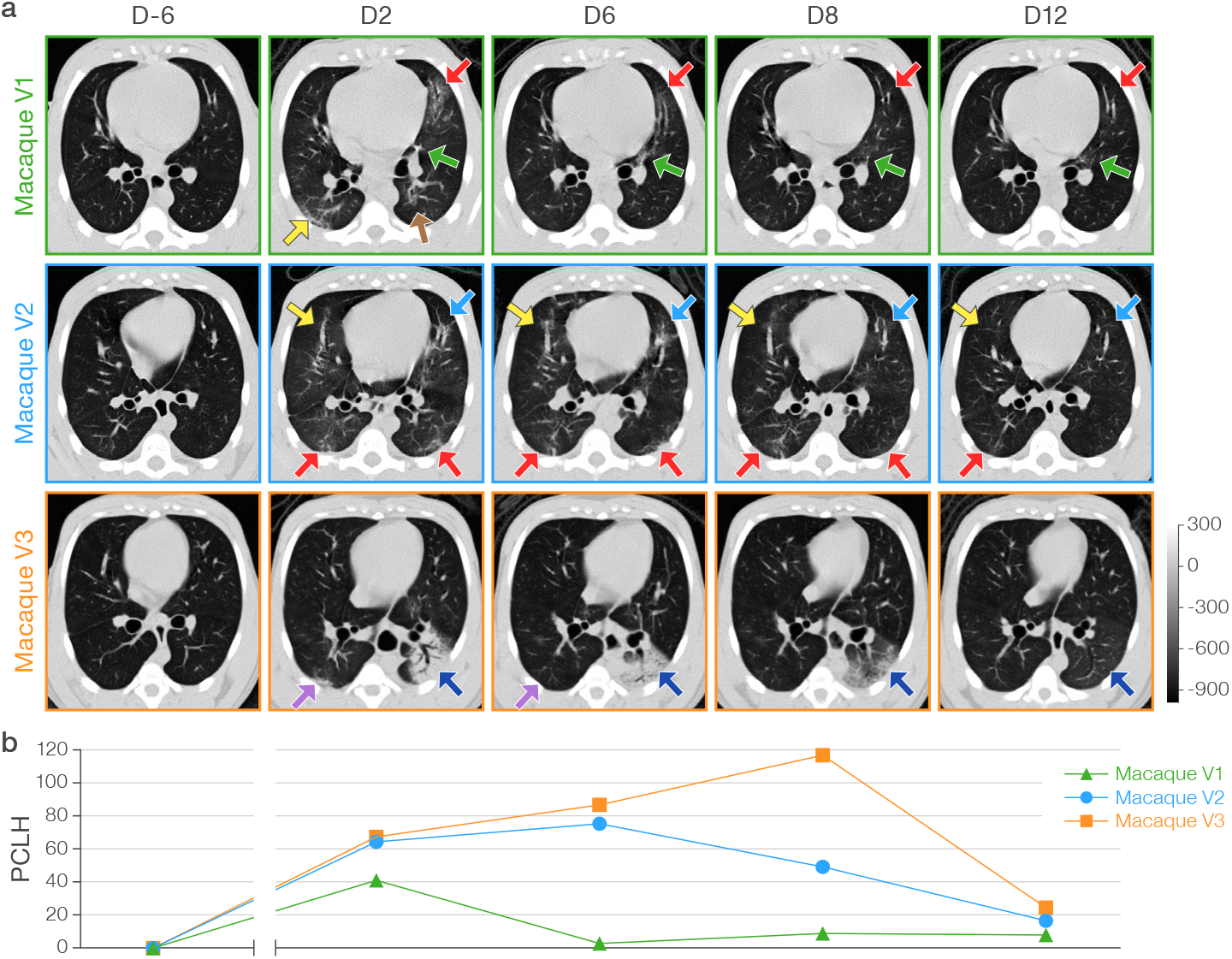
Qualitative and quantitative computed tomography (CT) analysis of macaque lungs. **a)** Representative axial CT images in three SARS-CoV-2-infected (V) macaques for each indicated study day (D). The grey scale represents radiodensity in Hounsfield units (HU). **b)** Percent change in volume of lung hyperdensity (PCLH) measured over time in the same SARS-CoV-2 inoculated macaques shown in a). Representative axial CT images and PCLH for all study days in both groups, including data for mock group (M) macaques, are shown in **Supplementary Figure 2.** In SARS-CoV-2-inoculated macaques only, selected representative CT images (axial, sagittal, and coronal views) with detailed radiological descriptions from pre-inoculation baseline to day 19) are shown in **Supplementary Figure 3.**

**Figure 5.**
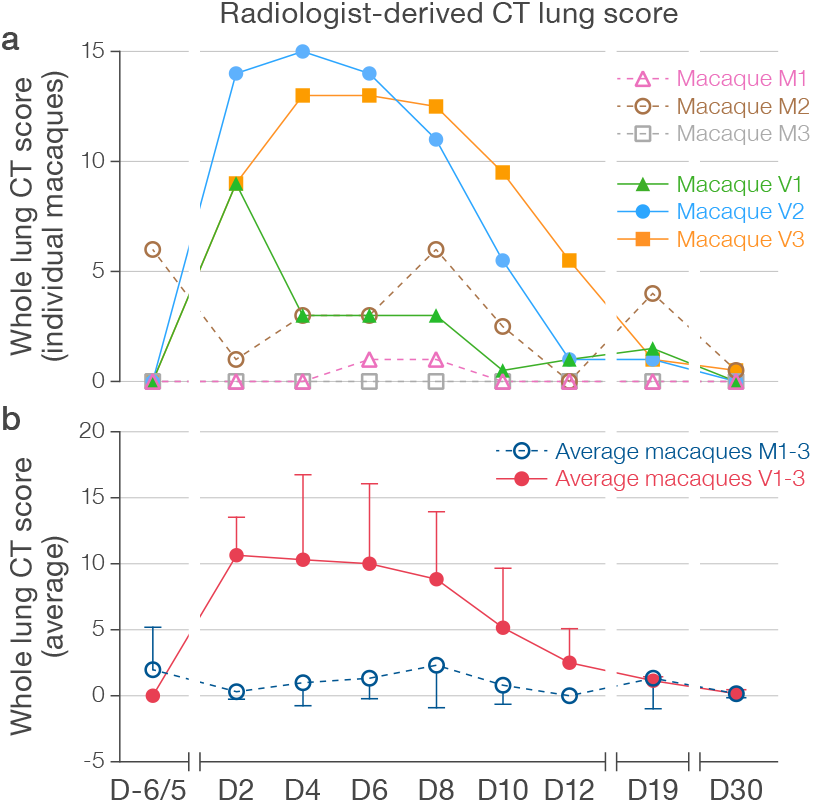
Radiologist-derived CT lung scores of macaque lungs. Averaged radiologist-derived CT scores of the entire lungs for individual macaques **(a)** and averaged for mock group (M) and virus group (V) macaques **(b)** over time.

Increased FDG uptake detected by PET (**Figure 6, Supplementary Figures 4–5**) corresponded well to the structural changes in the lungs observed by CT, and regional lymph node uptake was seen in all virus group macaques at D2. In macaque V1, FDG uptake decreased in the lungs at D6 but increased in mediastinal lymph nodes, and new FDG uptake was identified in the spleen. The two macaques (V2, V3) with persistent or progressive structural abnormalities on chest CT had variable changes in FDG uptake associated with the structural abnormalities in the lungs (some markedly increased, some improved) with an accompanying marked increase in FDG uptake in regional lymph nodes and spleens on D6. PET scan on D12 revealed normalization of previous areas of increased FDG uptake in the lung parenchyma in all three virus-exposed macaques, and persistent increased FDG update in regional lymph nodes and spleen. Mock-exposed macaques did not have similar increased FDG uptake with the exception of transient increased uptake in regional lymph nodes after mock-exposure in a single macaque (M1). Quantification of the SUV_max_ in selected regions of interest (ROI) in the lung, specific regional lymph nodes, and spleen mapped well to the qualitative findings in both mock-exposed and virus-exposed macaques **(Figure 7)**.

**Figure 6.**
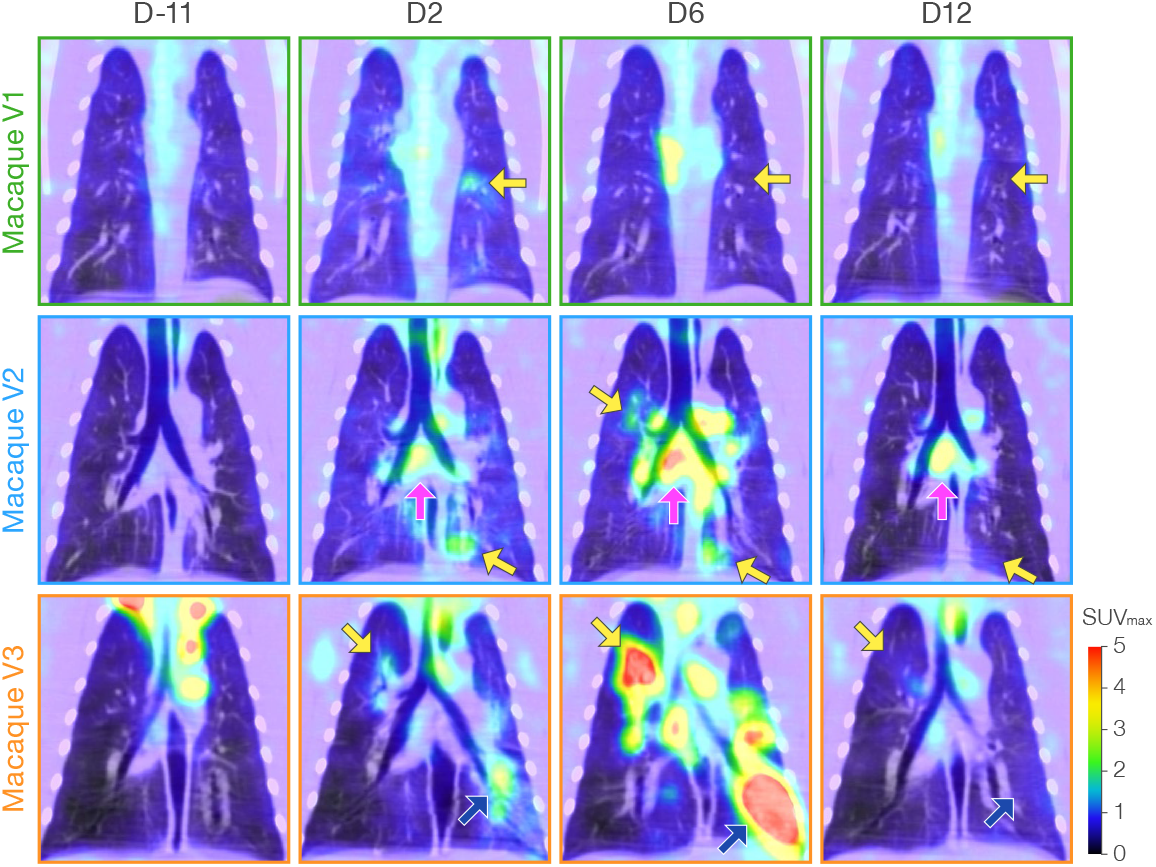
Qualitative positron emission tomography (PET) and FDG uptake analysis in macaque lung and regional lymph nodes. Representative coronal 2-deoxy-2-[^18^F]-fluoro-D-glucose (FDG) PET/CT images for each indicated study day (D) from pre-inoculation baseline to 12 days after inoculation. SUV_max_, maximum standardized FDG uptake value. Selected areas of increased FDG uptake are indicated in lung parenchyma (yellow arrows, blue arrows) and regional lymph nodes (pink arrows). All study days and data for mock group (M) macaques are shown in **Supplementary Figure 4**. Selected PET/CT images (axial, sagittal, and coronal views) including detailed radiological descriptions are shown in **Supplementary Figure 5**.

**Figure 7.**
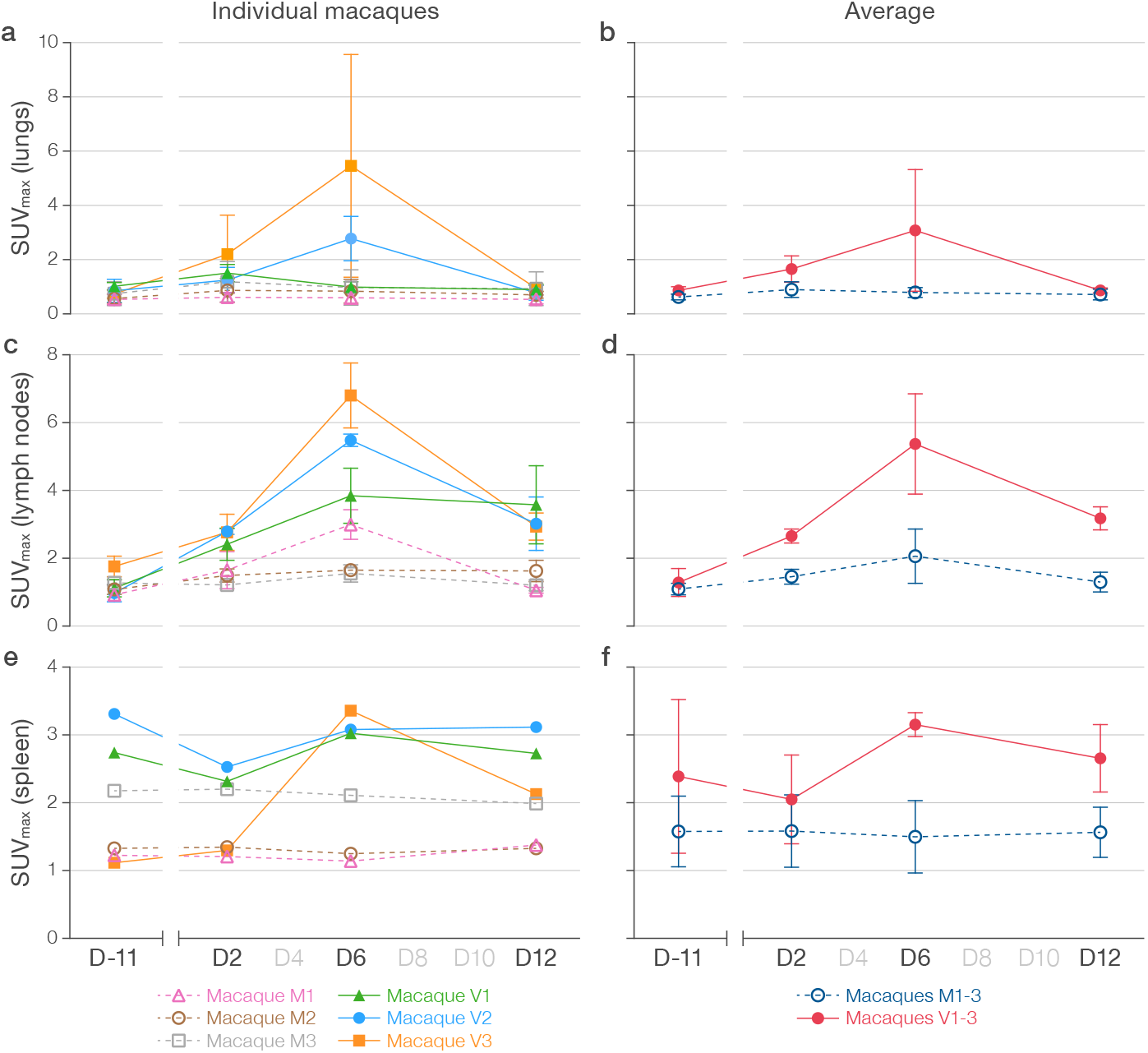
Quantitation of 2-deoxy-2-[^18^F]-fluoro-D-glucose (FDG) uptake in lung, lymph nodes, and spleen. **a)** SUV_max_ (mean FDG maximum standardized uptake values) for 3–5 selected lung regions of interest (ROIs) with high FDG uptake with tracking of identical regions of interests at all PET/CT timepoints in each macaque and **b)** averaged for mock group (M) and virus group (V) macaques. **c)** SUV_max_ for 2–3 selected lymph node ROIs with high FDG uptake in each macaque and **d)** averaged for (M) and (V) group macaques. **e)** SUV_max_ for spleen FDG uptake in each macaque and **f)** averaged for (M) and (V) group macaques.

CT images can be used for quantification of lung abnormalities using measures of volume or radiodensity, i.e., total lung volume (LV); average radiodensity in the total lung volume (LD); hyperdense volume (HV), a volume of lung in which radiodensity (Hounsfield units, HU) is above a pre-defined threshold; and average radiodensity in the hyperdense volume (hyperdensity, HD). Normalized changes from a pre-exposure baseline can be longitudinally measured as the percent change in the volume of lung hyperdensity (PCLH). Toward standardization across lung volumes, PCLH can be also be expressed as a percent of total lung volume (PCLH/LV). Increases in PCLH or PCLH/LV were not seen in the mock-exposed macaques over the entire study (**Figure 8a–d, Supplementary Figure 2).** However, post-exposure increases in PCLH or PCLH/LV were noted in all virus group macaques starting at D2, notably with heterogeneity in both the peak and duration of PCLH and PCLH/LV that corresponded well to longitudinal qualitative chest CT observations in individual virus group macaques (**Figure 4b, Figure 8a–d, Supplementary Figure 2).** Though both measures captured similar differences between groups, the within-group variability was unsurprisingly less with PCLH/LV versus PCLH. The virus group had significantly higher cumulative PCLH/LV over days 0–30 as summarized by the area under the curve (AUC_0–30_; *p*=0.01). The AUC_0–8_ for days 0–8 showed a similar trend (*p*=0.06), as did the AUC_0–8_ and AUC_0–30_ for the PCLH (*p*=0.06 and *p*=0.03, respectively).

**Figure 8.**
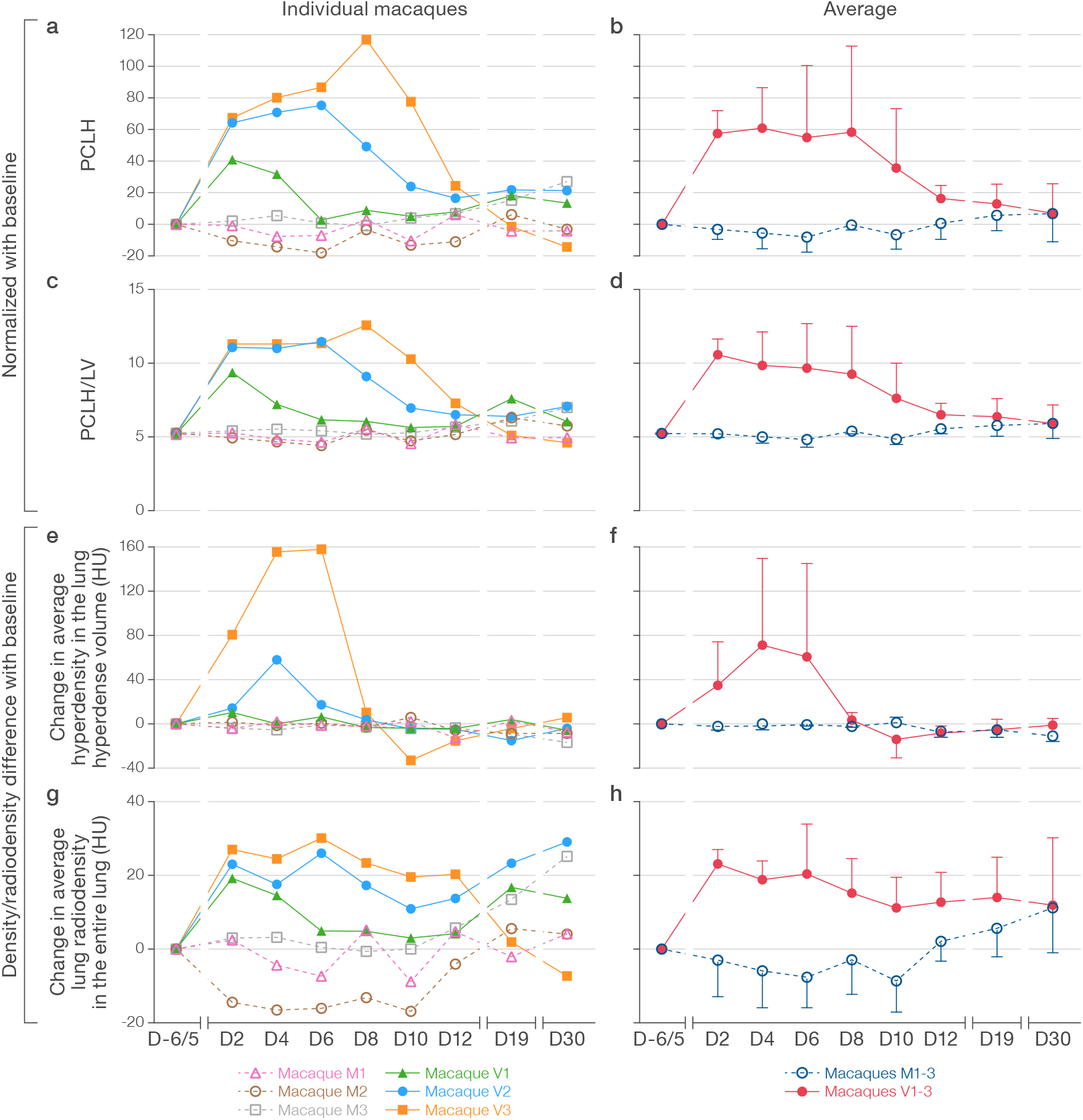
Quantitative analyses of volume and radiodensity of macaque lungs. **a–b)** Percent change in volume of lung hyperdensity (PCLH) measured over time for individual macaques (**a)** as in **Figure 4b** and averaged for mock group (M) and virus group (V) macaques **(b)**. **c–d)** PCLH standardized as percent of entire overall lung volumes over time (PCLH/LV) for individual macaques **(c)** and averaged for M and V macaques **(d)**. **e–f)** Change in the average lung densities (Hounsfield units [HU]) in the entire lung volumes over time for individual macaques **(e)** and averaged for M and V macaques **(f)**. **g–h)** Change in average lung hyperdensities (HU) in the lung hyperdense volume over time for individual macaques **(g)** and averaged for M and V macaques **(h)**.

A comparison of PCLH or PCLH/LV (**Figure 8a–d**) and absolute radiodensity (change in HD, change in LD) (**Figure 8e–h)** highlights similarities and differences windowed by these readouts that are particularly apparent as the CT abnormalities evolved in macaque V3. In this macaque, dense consolidation in the left lung reached peak radiodensity at D6, subsequently evolving toward a larger volume of less dense mixed consolidation and GGO at D8 **(Figure 4a).** This progression of findings is captured as an increase in PCLH **(Figure 8a)** and PCLH/LV **(Figure 8c)** from D6–8, but a sharp decline in HD **(Figure 8e)** over the same period.

Quantifiable changes in CT lung abnormalities, e.g., AUC of the PCLH/LV curve in an appropriately powered macaque study, could be used to objectively evaluate efficacy of candidate MCMs, including vaccines and therapeutics. Although the described crab-eating macaque model windows only mild to moderate radiographic disease, it captures heterogeneity in both severity and duration of disease; the readout can be similarly applied to any larger animal model of increased severity should they become available in the future. These objective measurements add to semi-quantitative scoring, which is potentially subject to observer bias in experimental settings (36–39); in our study, semi-quantitative findings include radiologist-derived CT lung scores (**Figure 5**) or mean SUV_max_ measured in operator-selected regions of interest (ROIs) on PET/CT scans (**Figure 7**).

Of interest, the disease heterogeneity captured by our imaging readouts is mirrored to some degree in thus-far limited measures of innate or adaptive immunity, namely in ELISA and neutralizing antibody titers and in longitudinal measurements of peripheral cytokines. Conclusions cannot be meaningfully drawn from the small numbers of measurements taken from only three macaques; nonetheless, between- and within-group differences in cytokines that have been identified as biomarkers of disease, disease severity, and disease outcome in humans (*30*, 40–44) are observed. Most notable in this regard is a remarkable concentration increase of cytokines associated with cytokine release syndrome (aka “cytokine storm”), such as C-X-C motif chemokine ligand 8 (CXCL8), interleukin (IL) 6, IL13, IL15, IL1 receptor antagonist (IL1RN), and tumor necrosis factor (TNF), starting around D6 in the macaque (V3) with the most significant CT and PET/CT abnormalities.

A key advantage of quantifiable CT chest imaging readout over serial euthanasia studies, in addition to potentially reduced experimental animal numbers, is the ability not only to evaluate between-group differences, but also to compare severity and duration of disease at higher resolution in single animals and even in isolated parenchymal areas sequentially. This approach can reduce the error inherent in cross-sectional sampling of individual animals at single timepoints. Imaging does, however, introduce its own experimental complexities and limitations. As we aimed to evaluate whether PCLH (or other CT imaging readouts presented in **Figure 8**) could be a useful quantitative readout for radiographic progression in the SARS-CoV-2 infected lung, we chose not to include irradiated inactivated SARS-CoV-2 in the mock inoculum to avoid antigen-induced inflammation and related radiographic changes. For similar reasons, and to avoid artificial dissemination of SARS-CoV-2, we specifically did not perform bronchoalveolar lavage (BAL) to obtain lung samples for downstream cellular, molecular, and virologic analysis (*45*, 46) and did not perform lung biopsies. The frequency of anesthesia and instrument availability pragmatically limit imaging to carefully chosen timepoints during a study. In particular, the extended time required to perform PET imaging resulted in logistical limitations of the number of macaques that could be included in the study. Finally, with complete resolution of radiographic abnormalities by the end of the study period, we opted not to euthanize these macaques to be able to perform a re-exposure study in the future. Thus, we cannot correlate radiographic with histopathologic findings.

Future studies should extend our initial findings in several directions. First, follow-up confirmation of these pilot results in this model of mild-moderate COVID-19 is needed to further establish quantifiable lung CT as a reliable disease readout and to forge imaging-pathologic correlates in macaques euthanized at peak radiographic abnormality. Confirmation should enable proof-of-concept evaluation of whether a candidate MCM will indeed significantly decrease peak or AUC of PCLH or PCLH/LV compared to untreated infected control macaques. Data from additional macaques will be used to confirm the sensitivity and relevance of the AUC_0–8_ and AUC_0–30_ for PCLH or PCLH/LV as robust measures of lung changes from CT evaluation.

In parallel, disease severity could possibly be increased in the crab-eating macaque model by optimizing delivery of SARS-CoV-2 to the most vulnerable lung (via aerosol or more distal bronchoscopic delivery), with the ultimate goal of using the CT-quantifiable volume or radiodensity readouts to model the sick hospitalized human.

Other groups are already evaluating NHPs of diverse species as possible COVID-19 models. In these models, serial chest CT imaging after intrabronchial instillation of SARS-CoV-2 could be used to establish a meaningful and quantifiable COVID-19-like disease readout that will enable objective evaluation of medical countermeasures and also a comparison of SARS-CoV-2-induced lung abnormalities in different NHP models.

## Methods

### Virus

Severe acute respiratory syndrome coronavirus 2 (SARS-CoV-2; *Nidovirales*: *Coronaviriridae*: *Sarbecovirus*) isolate 2019-nCoV/USA-WA1-A12/2020 was obtained from the US Centers for Disease Control and Prevention (CDC; Atlanta, GA, USA). A master virus stock (designated IRF_0394) was grown under high (biosafety level 3) containment conditions at the IRF-Frederick by inoculating grivet (*Chlorocebus aethiops*) Vero cells obtained from the American Type Culture Collection (ATCC; Manassas, VA, USA; #CCL-81) maintained in Dulbecco’s Modified Eagle Medium with L-glutamine (DMEM, Lonza, Walkersville, MD, USA) supplemented with 2% heat-inactivated fetal bovine serum (FBS; SAFC Biosciences, Lenexa, KS, USA) at 37°C in a humidified 5% CO_2_ atmosphere, harvested after 72 h, and quantified by plaque assay in Vero E6 cells (ATCC #CRL-1586) using a 2.5% Avicel overlay with a 0.2% crystal violet stain at 48 h following a previously published protocol (47). The genomic sequence of the IRF_394 master stock was determined experimentally by two independent amplification approaches: nonspecific DNA amplification (sequence-independent single primer amplification [SISPA]) as described previously (48) and the ARTIC protocol (49), which was designed to amplify overlapping regions of the SARS-CoV-2 reference genome (MN908947.3). Primer information and genomic alignment position are available at https://github.com/artic-network/artic-ncov2019/tree/master/primer_schemes/nCoV-2019/V1. PCR products were purified with the MinElute PCR Purification Kit (QIAgen, Valencia, CA, USA). Libraries were prepared with the SMARTer PrepX DNA Library Kit (Takara Bio, Mountain View, CA, USA), using the Apollo NGS library prep system (Takara Bio, Mountain View, CA, USA). Libraries were evaluated for quality using the Agilent 2200 TapeStation System (Agilent, Santa Clara, CA, USA). After quantification by qPCR with the KAPA SYBR FAST qPCR Kit (Roche, Pleasanton, CA, USA), libraries were diluted to 2 nM, and sequenced on a MiSeq (Illumina, San Diego, CA, USA). The genomic sequence of IRF_394 was found to be identical to the type sequence of SARS-CoV-2 isolate 2019-nCoV/USA-WA1-A12/2020 (GenBank MT020880.1), and IRF_394 was determined to be devoid of bacterial or viral contaminants.

### Animals

Six crab-eating (aka cynomolgus) macaques (*Macaca fascicularis* Raffles, 1821) of both sexes, 4–4.5 years old and weighing 3.17–4.62 kg (**Supplementary Table 1**), were obtained from Cambodia via Envigo Captive (Hayward, CA, USA) and housed at the US National Institutes of Health Animal Center (NIHAC; Dickerson, MD, USA) for 3 months. All female macaques were on depot medroxyprogesterone acetate (administered intramuscularly, 150 mg/ml) while at NIHAC for several months. The last dose administered was administered approximately one month prior to study start. The macaques were subsequently moved into the maximum (biosafety level 4 [BSL-4]) containment laboratory at the IRF-Frederick, a facility accredited by the Association for Assessment and Accreditation of Laboratory Animal Care International (AAALAC). Prior to facility entry, all macaques were serologically screened for herpes B virus, simian immunodeficiency virus (SIV), simian retrovirus, and simian T-lymphotropic virus (STLV) infection; all macaques tested negative. Macaques also tested negative multiple times for *Mycobacterium tuberculosis* infection. Once in containment, the macaques passed physical exams and routine bloodwork and were confirmed appropriate for study assignment by IRF-Frederick veterinarians. Experimental procedures for this study (protocol “SARS-CoV-2-NHP-064E-1”) were approved by the National Institute of Allergy and Infectious Diseases (NIAID), Division of Clinical Research (DCR), Animal Care and Use Committee (ACUC), and were in compliance with the Animal Welfare Act regulations, Public Health Service policy, and the *Guide for the Care and Use of Laboratory Animals 8^th^ Ed*. recommendations. The macaques were singly housed during the 2-week acclimatization to the maximum containment laboratory and the course of the study, and were provided with appropriate enrichment including, but not limited to, polished steel mirrors, durable toys, and food enrichment. Macaques were anesthetized in accordance with maximum containment standard operating procedures prior to all macaque manipulations, including virus exposure, sample collection, and medical imaging. Macaques were observed following anesthesia to ensure complete recovery. All work with NHPs was performed in accordance with the recommendations of the Weatherall Report.

### Macaque exposures

The macaques were split into 2 groups of 3 animals each (**Supplementary Table 1**). Mock group (M) macaques received 2 ml of DMEM + 2% heat-inactivated FBS into each bronchus by direct bilateral primary post-carinal intrabronchial instillation, followed by a 1-ml normal saline flush and then 5 ml air. Virus group (V) macaques were exposed the same way with each 2-ml instillate containing 9.13×10^5^ pfu/ml (i.e., a total exposure dose of 3.65×10^6^ pfu) of SARS-CoV-2 followed by 1-ml saline flush and then 5 ml air. All macaques were sedated prior to instillation. Prior to administering anesthesia, glycopyrrolate (0.06 mg/kg) was delivered intramuscularly to reduce saliva secretions. Next, each macaque received 10 mg/kg ketamine and then 35 μg/kg dexmedetomidine intramuscularly. To reverse anesthesia, 0.15 mg/kg atipamezole was administered intravenously. All macaques were evaluated daily for health and were periodically examined physically, including blood draws, and conjunctival (left and right), nasopharyngeal, oropharyngeal, and rectal swab collection. Stool and urine were also collected on each day swabs were collected. All swabs were collected in 1-ml universal virus transport (UVT) media (BD Biosciences, San Jose, CA, USA).

### Macaque scoring

Cage-side assessment scoring criteria (**Supplementary Table 4**) were modified from Chertow *et al.* (2016) to include clinical signs relevant to COVID-19 and respiratory rates of crab-eating macaques (*9, 50*, 51). In addition to cage-side observations, physical exam scoring criteria were implemented to assess clinical conditions on days when macaques were anesthetized (**Supplementary Table 4**). Cage-side and physical exam scoring criteria were developed in collaboration with National Primate Research Centers (NPRCs) to standardize disease assessment and compare disease outcomes between NHP models. Heart rate was not incorporated into the physical exam scores until D2 because heart rate score was determined as beats per minute over baseline. Baseline heart rate was determined as an average over three timepoints, D-11, D-5/D-6, and D0 (except for macaque V3, for which heart rate was not recorded prior to dexmedetomidine administration).

### Clinical analysis

Blood samples were collected in Greiner Bio-One Hematology K_3_EDTA Evacuated Tubes (Thermo Fisher Scientific). Complete blood counts, including leukocyte differentials and reticulocyte counts (CBC/Diff/Retic), were determined using the Procyte DX (IDEXX Laboratories, Westbrook, ME, USA). The Catalyst One analyzer (IDEXX Laboratories) was used for biochemical analyses of serum samples, which were collected in Greiner Bio-One VACUETTE Z Serum Sep Clot Activator Tubes (Thermo Fisher Scientific). Samples were run on both machines the day of collection shortly following collection such that they were not stored prior to analysis. On D30, an equipment issue with the Procyte DX required that samples from the virus-exposed group be stored at 4°C for about 30 h prior to analysis. For a list of all measured parameters and their values, see **Supplementary Table 3**.

### Image acquisition

Following sample collection and intubation, macaques were moved to chest computed tomography (CT) or whole-body positron emission tomography combined with CT (PET/CT) imaging. For imaging procedures, each macaque was anesthetized intramuscularly with 15 mg/kg ketamine following 0.06 mg/kg glycopyrrolate intramuscularly. Anesthesia was maintained using a constant rate intravenous infusion of propofol at 0.3 mg/kg/min (except on D-11 when propofol at 0.2 mg/kg/min was used). Macaques were placed on the scanner’s bed in a supine, head-out/feet-in position and connected to a ventilator to facilitate breath holds, and vital signs were monitored throughout the imaging process.

#### High resolution chest CT

Chest CT scans were performed using the 16-slice CT component of a Gemini TF 16 PET/CT (Philips Healthcare, Cleveland, OH, USA) or a Precedence SPECT/CT scanner (Philips Healthcare). Chest CT images were acquired in helical scan mode with the following parameter settings: ultra-high resolution, 140 kVp, 300 mAs/slice, 1 mm thickness, 0.5 mm increment, 0.688 mm pitch, collimation 16×0.75, and 0.75 s rotation. CT image reconstruction used a 512×512 matrix size for a 250-mm transverse field-of-view (FOV), leading to a pixel size of 0.488 mm. Two CT images were produced: one with the standard “B” filter and one with the bone-enhanced “D” filter. No contrast agent was administered. Each macaque underwent a 15–20-s breath-hold during acquisition. The pressure for the breath-hold was maintained at 150 mmH_2_O.

#### Whole-body PET/CT

Whole-body PET/CT scans were performed using a Gemini TF 16 TF PET/CT scanner. Radiotracer (2-deoxy-2-[^18^F]-fluoro-D-glucose; FDG) was injected intravenously (0.5 mCi/kg FDG, up to 4.0 mCi/scan) and the time of injection was recorded. After high-resolution chest CT imaging with breath-hold session (≈5 min), whole-body CT images were acquired (≈5 min) in helical scan mode with the following parameter settings: high resolution, 140 kVp, 250 mAs/slice, 3 mm thickness, 1.5 mm increment, 0.688 mm pitch, collimation 16×0.75, and 0.5 s rotation. Two CT images were reconstructed from the raw data. An initial CT image was reconstructed into a 600-mm diameter field-of-view (FOV), resulting in a pixel size of 1.17 mm and a slice spacing of 1.5 mm. This CT image was used to create an attenuation map needed to correct the PET images for photon attenuation. Raw CT data were reconstructed a second time into diagnostic quality CT images by reducing the FOV size to 250 mm, resulting in a pixel size of 0.488 mm with 1 mm slice thickness. One CT image was produced with the standard (“B”) filter. No contrast agent was administered, and the macaques breathed freely during the scan. Following whole-body CT scanning, whole-body PET scans covering the macaques’ bodies from the top of the head to the middle of the thighs was performed after a 60-min delay. Depending on the size of the NHP, 6 or 7 bed positions (with 50% overlap) were needed for this scan range. Dwell time per bed position was 3 min, resulting in a total duration of 18 or 21 min/scan. PET data were reconstructed into a set of either 300- or 342-image slices with 128×128 2-mm-cubic voxels. To ensure quantitative accuracy, all reconstructed PET images were corrected during scans for radioactive decay, uniformity, random coincidences, and attenuation and scattering of PET radiation *in situ*. Lastly, whole-body CT imaging with iopamidol (600 mg iodine/kg) intravenous contrast material were acquired at D-11 and the terminal D30 scans. Each macaque underwent a 30–50-s breath-hold during acquisition. The pressure for the breath-hold was maintained as 150 mmH_2_O. After completion of imaging, macaques were returned to the clinical team for subsequent procedures.

### Image evaluation

#### Chest CT evaluation

An adapted semi-quantitative scoring system based on a previously published method (52) was used to quantitatively estimate the pulmonary involvement of lung abnormalities on the extent of parenchymal lung disease in each lung lobe. The sums of the lobar scores were used to generate total lung summary scores. CT scores ranged from 0–5 for each lobe (right upper, right middle, right lower, right accessory, left upper, left middle, and left lower). Lobes were scored as: 0 = no disease involvement, 1 ≤5%, 2 = 5– 24%, 3 = 25–49%, 4 = 50–74%, 5 ≥75%, with a maximum total lung score of 35. Score increments of 0.5 were used to indicate improvement (or worsening) in radiodensity when changes in volume were insufficient to change score category. Additional data points included cohort (macaques M1–3 and V1–3), scan date and study day, and number of “lesions”. Types of infiltrates (GGOs, paving, consolidation, and organizing pneumonia) per lobe were given individual 0–5 scores, and the overall predominant type of infiltrate per each lobe and each subject scan were recorded. The summary scores per each lobe and whole lung were used as markers of disease progression. Image analysis was performed by a board-certified radiologist and a research fellow using MIM software version 6.9 (Cleveland, OH, USA).

To quantify CT data, the lung field was segmented using a region-growing implementation (MIM software). Entire lung volumes (LV) were measured at each time point (n: at time point n; b: at baseline). A histogram analysis was performed on the voxel intensities (radiodensity in Hounsfield units [HU]) within the segmented lung. Percent change in the volume of hyperdense lung tissue was determined as described previously (53). Briefly a threshold value was determined for each subject, based on a 5% cutoff in the upper tail of the histogram of lung tissue from the baseline CT scan. Due to an inability to keep the lungs of the virus group macaques inflated to approximately the same volumes over time, a correction was applied to the 5% PCLH threshold as previously described (54).

As more abnormalities (e.g., GGOs, consolidation) appear during the disease process, a larger volume of tissue will have higher HU values and PCLH as percent change in lung hyperdense volume (HV) from baseline can be expressed as [(HV_n_-HV_b_)/HV_b_]*100. Then PCLH/LV can be expressed as [V_n_/LV_n_]*100. Change in average lung radiodensity (LD) in the entire lung volume can be [LD_n_-LD_b_]. Change in average hyperdensity in hyperdense volume can be expressed as [HD_n_-HD_b_]. Then, PCLH as percent change in lung hyperdense volume, PCLH as percent of lung volume (PCLH/LV), change in the average radiodensity in the entire lung and change in the averaged hyperdensity in the hyperdense volume were graphed with disease progress. To visualize CT abnormalities in three dimensions (3D), a volume rendering technique was used to create videos. In brief, the lungs and airways were extracted form chest CT images using MIM software. A region growing algorithm was used to segment different classes including normal lung tissue, vessels, airways, and “lesions” with multiple seeds at specific locations to achieve realistic segmentations. 3D volume renderings of the segmentations were generated and animated rotations exhibiting the location and extent of the abnormalities were produced using 3D Slicer 4 software version 4.10.2 (55).

#### Whole-body PET/CT evaluation

Analysis of imaging data was performed using MIM software. Whole-body CT and PET scans for a given scanning session were co-registered. Regions of interest (ROIs) were placed manually on the PET scans, and location determined on the co-registered CT scans. These regions included specific lung abnormalities when present, left and right lung when no specific abnormalities were present, mediastinal and hilar lymph nodes, and spleen. Once ROIs were placed, mean FDG maximum standardized uptake values (SUV_max_) were measured from corrected PET images and averaged SUV_max_ values were graphed longitudinally.

The quantitative analysis was correlated with a qualitative evaluation of CT and PET/CT lung pathology over time, performed by a board-certified radiologist.

### RT-qPCR

RT-qPCR analysis was performed to determine presence of SARS-CoV-2 RNAs in collected specimens. Samples were frozen at −80°C in TRIzol LS (Thermo Fisher Scientific, Wilmington, DE, USA) and thawed on ice. 100 μl of sample were added to 5PRIME Phase Lock tubes (Quantabio, Beverly, MA, USA) followed by addition of 20 μl of chloroform/tube (Sigma-Aldrich, St. Louis, MO, USA) and inversion by hand 10 times. Phase Lock tubes with sample and chloroform were centrifuged at 10,000 x *g* for 1 min at 4°C. Following centrifugation, aqueous phases were removed (≈55 μl/tube) and deposited into clean 1.5-ml Eppendorf tubes. 70% ethanol was subsequently added to each tube at a 1:1 ratio (55 μl ethanol), inverted 10 times by hand, and briefly centrifuged. The ethanol/aqueous solution containing extracted RNA was used as input for purification and isolation using the PureLink RNA Mini Kit (Thermo Fisher Scientific) following the manufacturer’s instructions. RNA was eluted in 30 μl of water. RT-qPCR was performed using the SuperScript III Platinum One-Step qRT-PCR Kit (Thermo Fisher Scientific) following the manufacturer’s instructions with the following changes: reagent volumes were halved, resulting in 25-μl final reaction volumes. The N1 assay supplied with the 2019-nCoV CDC qPCR Probe Assay (Integrated DNA Technologies [IDT], Coralville, IA, USA) was used (1 μl/reaction) in lieu of individual primers and probes. 2 μl of extracted RNA or 2019-nCoV_N_Positive_Control (IDT) were used in each reaction. Samples and controls were run in technical triplicates on a CFX96 Touch Real-Time PCR Detection System (Biorad, Hercules, CA, USA) following the manufacturer’s recommendations; a 50°C, 15-min reverse-transcriptase step was used for first strand cDNA synthesis followed by 95°C, 2-min to inactivate the reverse transcriptase, followed by 45 cycles of 95°C, 15 s, 60°C, 30 s.

### Serology

Serum samples were collected in Greiner Bio-One VACUETTE Z Serum Sep Clot Activator Tubes and frozen at −80°C. Before removal from the maximum containment laboratory, virus in samples was inactivated using a cobalt irradiation source with a target dose of 50 kGy (JLS 484R-2 Cobalt-60 [^60^Co] Irradiator, JLShephard & Associates) following standard inactivation protocols. Serum samples were subsequently heat-inactivated at 56°C for 30 min prior to antibody screening. To determine IgG titers, Immulon 2HB 96-well plates (Thermo Fisher Scientific) were coated with recombinantly expressed SARS-CoV-2 spike S1 subunit (Sino Biological, Wayne, PA, USA) at 0.1 μg/well in 50 μl/well overnight at 4°C. Plates were washed three times with phosphate-buffered saline (PBS) + 0.1% Tween20 (PBS_T; Sigma-Aldrich) and blocked with ELISA diluent (5% nonfat milk [LabScientific, Danvers, MA, USA] in PBS-T) for 1 h at 37°C. Serum samples were serially diluted 1:2 in a dilution block (1:50 to 1:6,400) with ELISA diluent. After blocking, plates were washed three times with PBS-T and 100 μl/well of diluted sample were transferred to the plate. Samples were incubated for 1 h at 37°C. Plates were then washed three times with PBS-T, and 100 μl of goat anti-human IgG Fc specific (Jackson ImmunoResearch, West Grove, PA, USA) secondary antibody conjugated to horseradish peroxidase (HRP; diluted 1:20,000 in ELISA diluent) were added to each well. Samples were incubated for 1 h at 37°C and finally washed five times with PBS-T. Plates were developed by adding 100 μl of TMB substrate (Thermo Fisher Scientific) at room temperature for 10 min in the dark. Development was stopped by the addition of 100 μl of Stop Solution (Thermo Fisher Scientific). Plates were read at 450 nm with a correction wavelength of 650 nm using a Spectramax Plus 384 (Molecular Devices, San Jose, CA, USA) within 30 min of stopping the reaction. Reciprocal endpoint titers were determined in GraphPad software version 8.4.2 (Prism, La Jolla, CA, USA), using a sigmoidal 4 parameter-logistic fit curve. Endpoint titers were calculated at the point when the curve crossed the ELISA cutoff value. For the S1 subunit IgG ELISA, the cutoff value was determined to be an optical density (OD) of 0.19077, which was determined from control sera collected from twenty-five NHPs sampled prior to the known emergence of SARS-CoV-2. Data are presented as the mean and standard deviation of two independent ELISA runs.

### Fluorescence neutralization assay

All assays were run with irradiated and heat-inactivated sera. Irradiation and heat-inactivation were performed as described above. Vero E6 cells were seeded at 3×10^4^ in 100 μl DMEM+10% heat-inactivated FBS in 96-well Operetta plates (Greiner Bio-One, Monroe, NC, USA). The following day, a series of six-point dilutions, each 1:2, were performed in duplicate starting with a dilution of 1:20 (1:20, 1:40, 1:60, etc.) in 96-well 1.2-ml cluster tubes (Corning Inc, Corning, NY, USA). Then, stock SARS-CoV-2 virus was diluted in serum free media and was added to the sera in each cluster tube at a multiplicity of infection (MOI) of 0.5 using a liquidator, doubling the total volume in each well and further diluting sera 1:2. Thus, the final starting dilution was 1:40. The sera/virus mixtures were then mixed by pipetting up and down with the liquidator. Cluster tubes were next incubated for 1 h at 37°C/5% CO_2_. Following incubation, 100 μl/well of each virus/serum mixture were transferred to the Operetta plates from the cluster tubes to yield final volumes of 200 μ/well (100 μl cell seeding media plus 100 μl virus/serum mixture). Each set of cluster tubes provided enough material for each sample to be run in duplicate in 2 plates, yielding a total of 4 replicates per sample. The virus/serum mixtures were incubated on plates for 24 h at 37°C/5% CO_2_. Subsequently, plates were fixed with 20% neutral buffered formalin (Thermo Fisher Scientific) for 24 h at 4°C. Next, plates were washed with PBS (Thermo Fisher Scientific) and then blocked with 3% bovine serum albumin (BSA) in 1X PBS for 30 min on a rocker. Staining followed, first with the primary antibody and then the secondary antibody. The primary antibody was SARS-CoV-2 Nucleoprotein/NP Antibody, Rabbit mAb (Sino Biological, Chesterbrook, PA, USA) prepared at 1:8,000 in blocking buffer at room temperature. Plates were incubated with primary antibody for 60 min on a rocker. The secondary antibody was goat α-rabbit IgG (H+L), Alexa Fluor 594 Conjugate (Thermo Fisher Scientific) prepared at 1:2,500 in 1X PBS. Plates were incubated with secondary antibody at room temperature for 30 min on a rocker and in the dark. Between the blocking, primary, and secondary steps, plates were washed three times with 1X PBS. Finally, plates were read on a Operetta High-Content Analysis System (PerkinElmer, Waltham, MA, USA) with at least 4 fields of view of>1,000 cells each. Data were analyzed using Harmony software (PerkinElmer). Half-maximal neutralization titers (NT_50_) were calculated by averaging the fluorescence intensity in virus control wells and dividing by two. The fluorescence intensity of a sample at each dilution was compared to the NT_50_ values, and the lowest dilution that is equal to or less than the NT_50_ value was recorded.

### Cytokine analysis

The MILLIPLEX MAP Non-Human Primate Cytokine Magnetic Bead Panel - Premixed 23 Plex – Immunology Multiplex Assay (Millipore, Burlington, MA, USA; #PCYTMG-40K-PX23) was performed on collected plasma samples following the manufacturer’s instructions. All reagents were warmed to room temperature prior to addition to assay wells. Quality controls and standards were reconstituted with 250 μl of deionized water and allowed to sit for 10 min prior to use. A four-fold seven-point standard curve was generated by diluting the concentrated stock with assay buffer for each point. Assay buffer alone served as the blank. The plate was first washed with 200 μl of assay buffer and incubated on an orbital shaker at 800 rpm at room temperature for 10 min. Assay buffer was decanted, and plates were inverted on absorbent paper to remove any excess buffer. 25 μl of serum matrix were added to background, control, and standard wells. 25 μl of assay buffer was added to sample wells. 25 μl of samples/standards/controls were added to the appropriate wells. Beads were resuspended by vortexing and 25 μl were of vortexed beads were added to each well. Each plate was sealed and incubated overnight at 4°C on an orbital shaker at 500 rpm. The next day, plates were washed twice using a hand-held magnetic plate holder with 200 μl of wash buffer per well and decanted as previously described. Plates were then incubated on an orbital shaker at 500 rpm at room temperature for 1 h with 25 μl of detection antibody. After incubation, 25 μl of kit-provided streptavidin-phycoerythrin were added directly to each well and incubated at room temperature on an orbital shaker at 500 rpm for 30 min. Plates were washed twice and 150 μl of sheath fluid was added to each well. Plates were read on a Flexmap 3D reader (Luminex, Chicago, IL, USA) within 24 h of completion following assay instructions. The data was exported to Bio-Results Generator version 3.0 and Bio-Plex Manager software version 6.2 (BioRad). Results were graphed using GraphPad software version 8.4.2.

### Statistical Analysis

Area under the curve (AUC) summaries were calculated using the trapezoidal rule, and compared using Welch’s t-tests, using R version 3.6.3.

### Data Availability

Data from this study were made available publicly as soon as they became available at https://openresearch.labkey.com/project/Coven/COVID-001/begin.view to inform the ongoing COVID-19 outbreak response without delay following practices we initiated for NHP studies of Zika virus pathogenesis in 2016 (56).

## Acknowledgements

We thank all the staff of the NIH/NIAID/DCR/Integrated Research Facility at Fort Detrick who supported this study, in particular Karlton Churchwell, David Drawbaugh, Kyra Hadley, Zachary Hubble, Nicolette Schuko, and Colin Waters (Clinical Core), Kurt Cooper and Dan Ragland (Comparative Medicine), Sean Bartlinski (Data Management), Elena N. Postnikova, Robin Gross, Shuiqing Yu, Lindsay Marron, Steve Mazur, Saurabh Dixit, Heema Sharma, and Huanying Zhou (Antibody Screening), Jurgen Seidel (Imaging), Rebecca Bernbaum, Blake Davis, and Erika Maynor (Immunology), Louis Huzella (Pathology), as well as Yu Cong and Travis K. Warren. We are grateful to Chelsea Crooks, Amelia K. Haj, Anna S. Heffron, Joseph Lalli, Trent M. Prall, and all other members of the Coven Consortium at University of Wisconsin for critical input into study design, study planning and support, and real-time data posting support. We also would like to thank Daniel S. Chertow (NIH/NIAID/DCR/Clinical Center) for crucial guidance on intrabronchial instillation, and Nathan W. Finch for his insight particularly with respect to our radiological findings and analyses.

This work was supported in part through Laulima Government Solutions, LLC prime contract with the US National Institute of Allergy and Infectious Diseases (NIAID) under Contract No. HHSN272201800013C (R.B., T.K.C., J.L., K.S., G.K., P.S., C.B., R.A., J.R.K., T.B., M.G.L., J.W.). C.L.F., J.H.L., and J.H.K. performed this work as employees of Tunnell Government Services (TGS), a subcontractor of Laulima Government Solutions, LLC under Contract No. HHSN272201800013C. This work was also supported in part with federal funds from the National Cancer Institute (NCI), National Institutes of Health (NIH), under Contract No. HHSN261200800001E to I.C. and J.S., who were supported by the Clinical Monitoring Research Program Directorate, Frederick National Lab for Cancer Research, sponsored by NCI. The views and conclusions contained in this document are those of the authors and should not be interpreted as necessarily representing the official policies, either expressed or implied, of the US Department of Health and Human Services or of the institutions and companies affiliated with the authors. The study protocol was reviewed and approved by the NIH/NIAID/DCR/Integrated Research Facility at Fort Detrick, Frederick, MD, USA Animal Care and Use Committee in compliance with all applicable federal regulations governing the protection of animals and research.

## Author contributions

C.L.F., I.C., J.H.L., T.K.C., J.R.K., M.C.St.C., M.G.L., R.F.J., K.M.B., M.R., C.S., T.C.F., D.H.O’C., and J.H.K. contributed to the study conception and design. C.L.F., J.H.L., R.B., T.K.C., J.L., K.S., P.J.S., G.K., C.B., P.A.L., R.A., B.B., N.D.P., J.R.K., T.B., M.C.St.C. contributed to study performance, and sample and data collection. C.L.F., I.C., J.H.L., R.B., T.K.C., J.L., J.S., P.S., C.B., N.A., M.C., T.C.F., P.A.L., B.B., N.D.P., J.R.K., J.R.K., M.C.M., I.M.F., G.P., J.W., T.C.F, D.H.O’C., and J.H.K. contributed to data analyses, interpretation, and writing. All authors read and approved the final manuscript.

**Videos 1–3 | Qualitative computed tomography (CT) analysis of macaque lungs.** 3D rendering of the lungs of macaque V1 at D2 **(Video 1)**, V2 at D4 **(Video 2)**, and V3 at D4 **(Video 3)** after SARS-CoV-2 exposure. Blue: airways; gray: normal lung; red: vessels; yellow: imaging abnormalities.

## Supplementary Data

**Supplementary Figure 1 | Cytokines.** Cytokien concentration changes measured for individual macaques and averaged for mock group (M) and virus group (V) macaques. Data are represented on logarithmic scales. For graphic representation, values of “0” or “not detected” (below the level of assay sensitivity) were automatically assigned a value = 1.

**Supplementary Figure 2 | Qualitative and quantitative computed tomography (CT) analysis of macaque lungs. a)** Representative axial CT images in three SARS-CoV-2-infected (V) macaques and mock-inoculated (M) macaques for all indicated study days (D). The grey scale represents radiodensity in Hounsfield units (HU). Selected CT images (axial, sagittal, and coronal views) from the SARS-CoV-2 infected macaques with detailed radiological descriptions are shown in **Supplementary Figure 3**. Colored arrows in **Figure 4** and this figure represent regions of interest that are further detailed in figure legends for **Supplementary Figure 3**. **b)** Percent change in lung hyperdensity (PCLH) measured over time in the same macaques shown in **Figure 4**, including also here the mock-infected individual macaques and all study days.

**Supplementary Figure 3 | Qualitative computed tomography (CT) analysis of macaque lungs.**

**Supplementary Figure 4 | Qualitative positron emission tomography (PET) and PET/CT analysis of macaque lungs.** Representative coronal (left panels) and axial (right panels) 2-deoxy-2-[^18^F]-fluoro-D-glucose (FDG) PET/CT images for each indicated study day (D). SUV_max_, mean FDG maximum standardized uptake values. Selected areas of increased FDG uptake are highlighted in the lung parenchyma (yellow arrows) and regional lymph nodes (pink arrows). Selected merged PET/CT images (axial, sagittal, and coronal views) with detailed radiological descriptions of these areas of interest are shown in **Supplementary Figure 5**.

**Supplementary Figure 5 | Qualitative positron emission tomography (PET) and PET/CT analysis of macaque lungs.**

**Supplementary Table 1 | Crab-eating macaque (*Macaca fascicularis* Raffles, 1821) information.**

**Supplementary Table 2 | Macaque physical condition/clinical scoring results.**

**Supplementary Table 3 | Complete blood cell count (CBC/Diff/Retic) and serum chemistries.**

**Supplementary Table 4 |Macaque clinical scoring guide.**

