## Supplementary figures and images for "Characteristic and quantifiable COVID-19-like abnormalities in CT- and PET/CT-imaged lungs of SARS-CoV-2-infected crab-eating macaques (*Macaca fascicularis*)"

### Supplemental Figure 1

#### Supplementary Figure 1

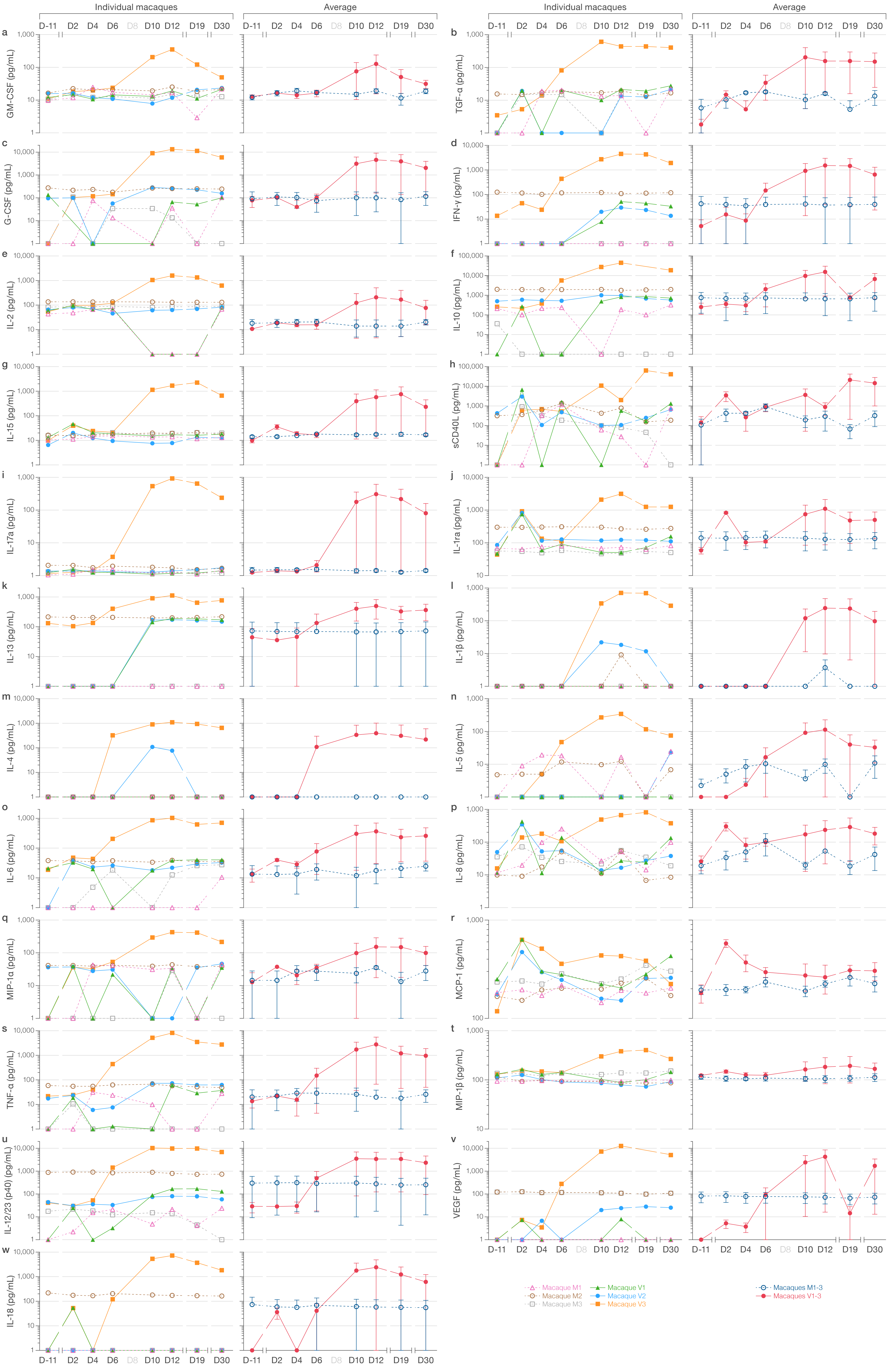

### Supplemental Figure 2

#### Supplementary Figure 2

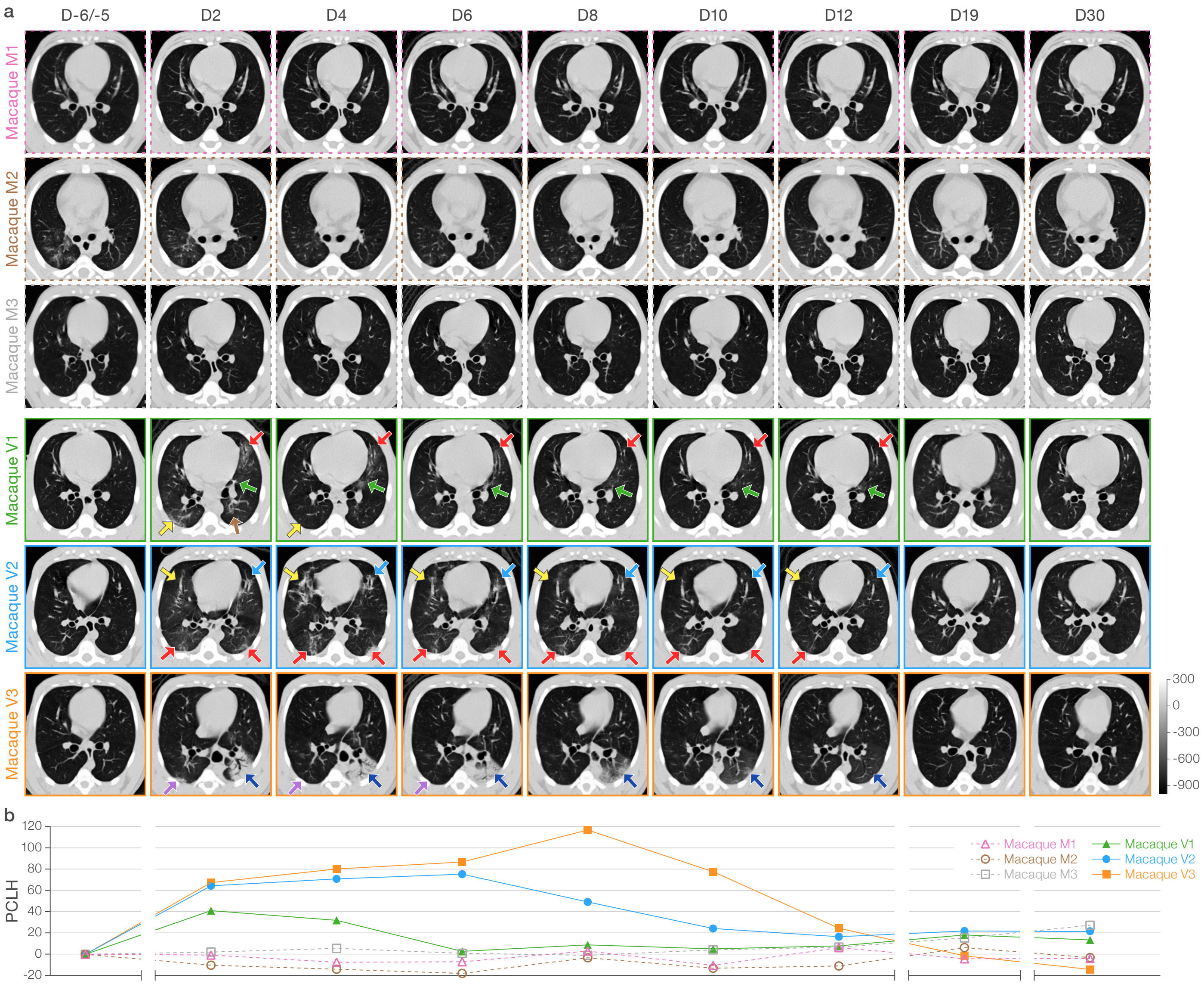

### Supplemental Figure 4

Supplementary Figure 4

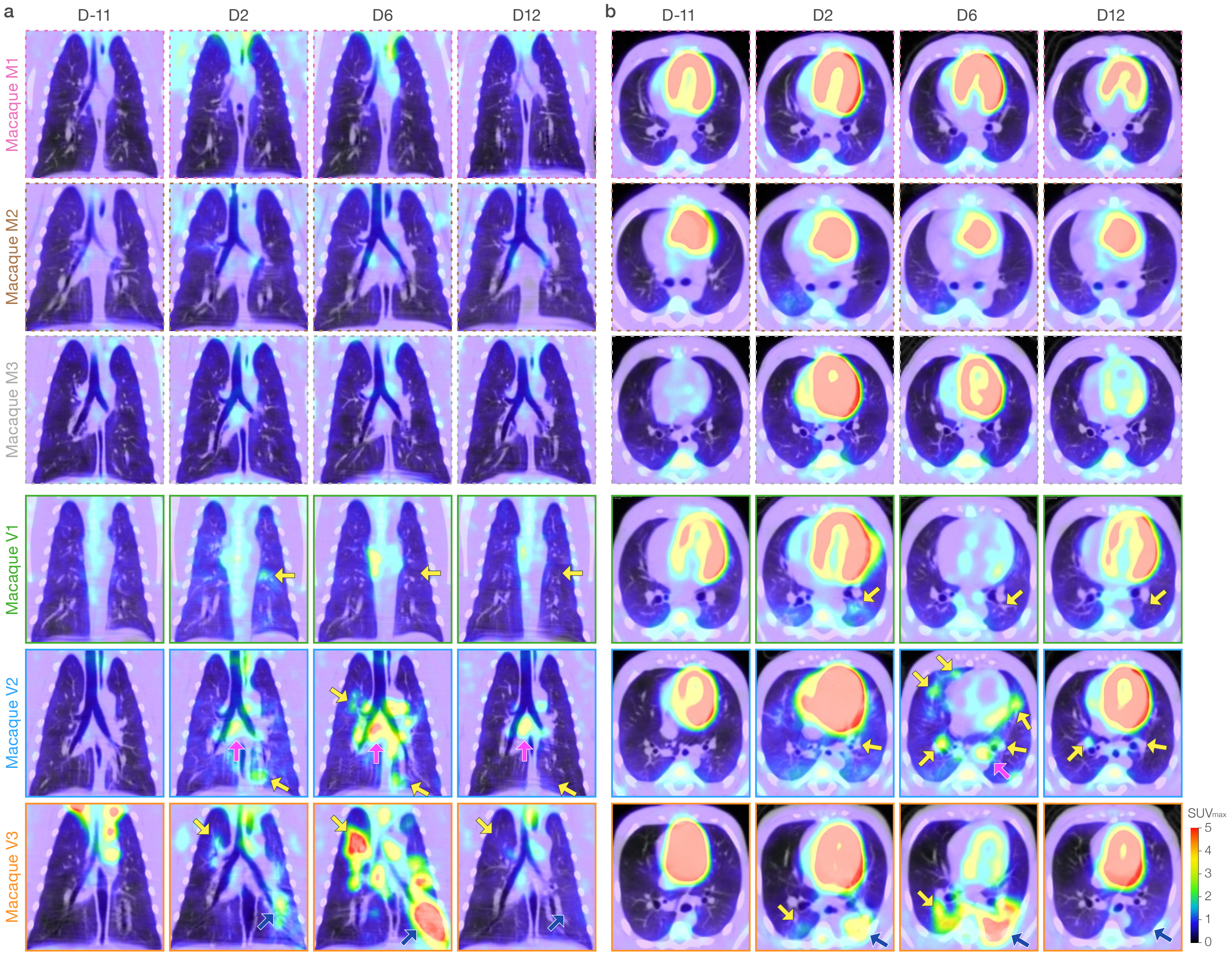
