## Supplemental Figure 3 for "Characteristic and quantifiable COVID-19-like abnormalities in CT- and PET/CT-imaged lungs of SARS-CoV-2-infected crab-eating macaques (*Macaca fascicularis*)"

Supplementary Figure 3

### Macaque V1

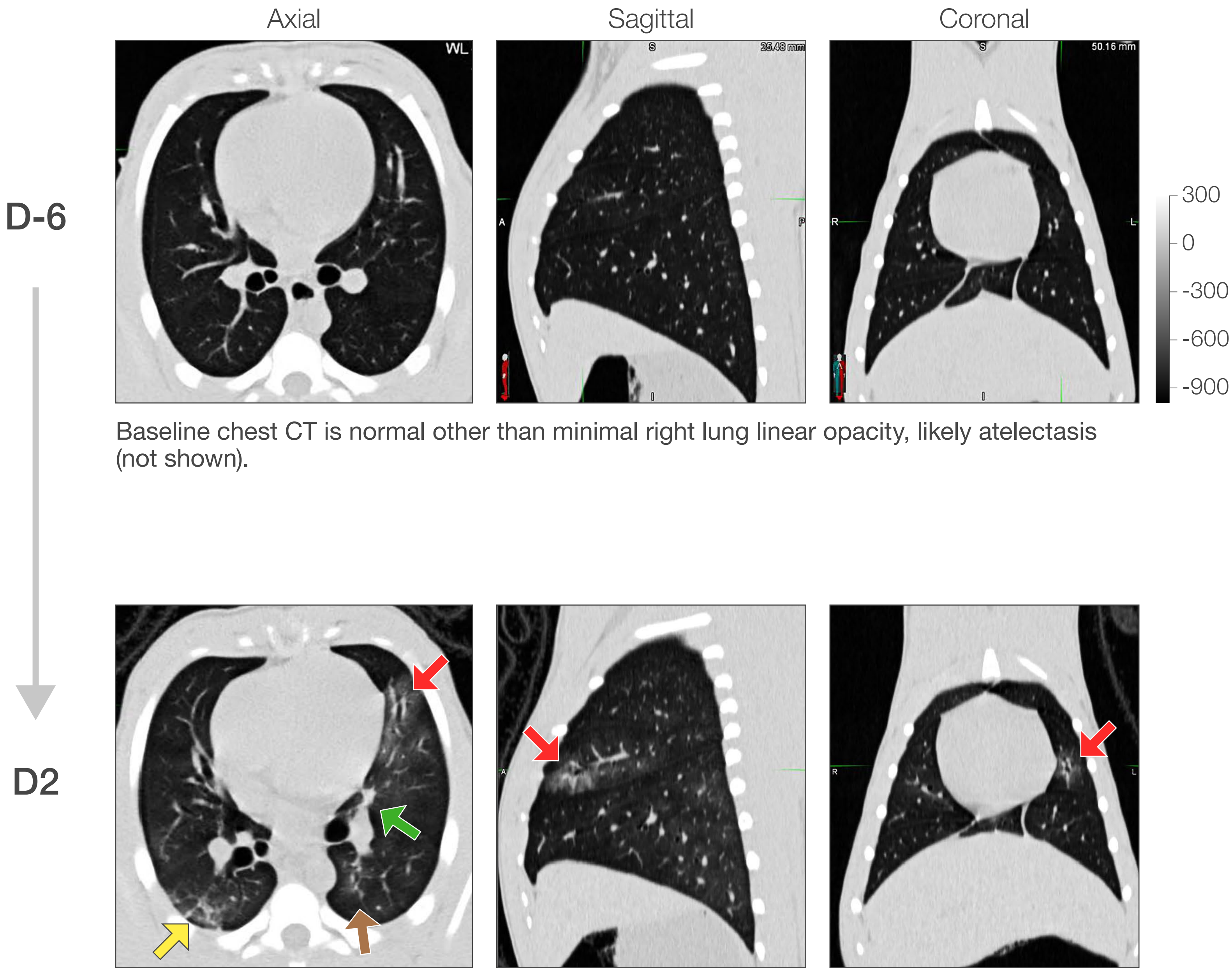


Baseline chest CT is normal other than minimal right lung linear opacity, likely atelectasis (not shown).

Compared to baseline, chest CT scan on D2 shows new bilateral, multilobar abnormalities, including patchy ground-glass opacity (GGO) surrounding peri-bronchial thickening (left middle lobe, red arrow), a small focus of consolidation (left middle lobe, green arrow), GGO with reticular infiltrate (posterior right lower lobe, yellow arrow) and right lower lobe subpleural bands (not shown). These infiltrates were FDG-avid by PET/CT scan (not shown). Subtle GGO (posterior left lower lobe, brown arrow).

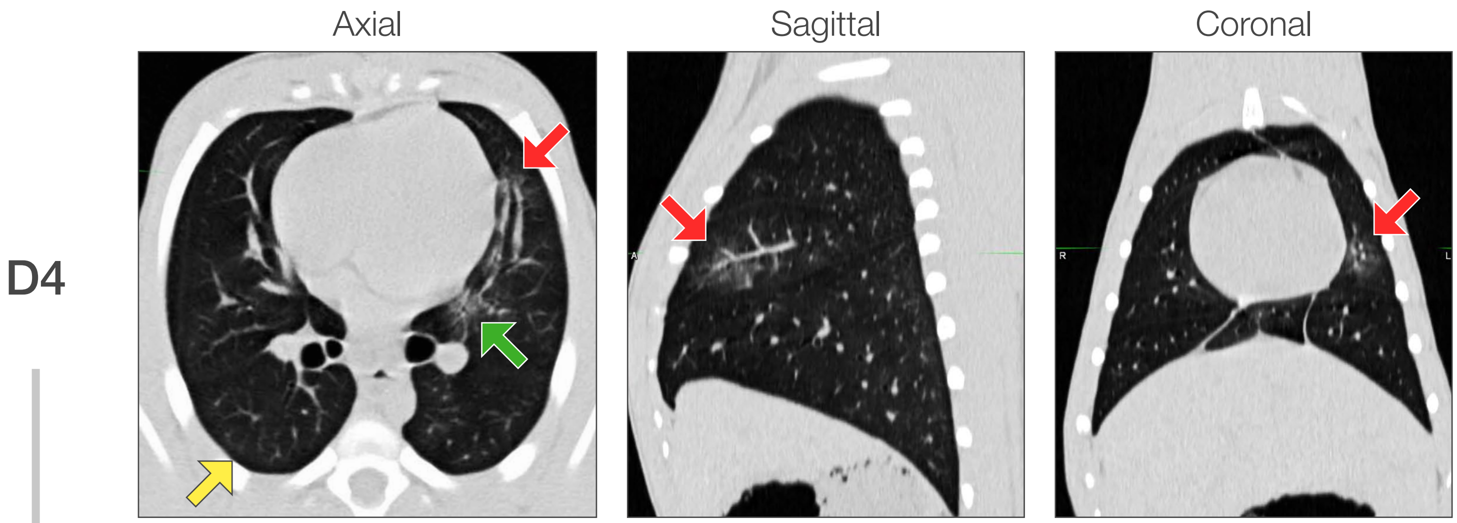


Compared to D2, chest CT on D4 showed resolution of the right lower lobe posterior infiltrate (yellow arrow) and sub-pleural bands (not shown), improvement in the radiodensity of the GGO and peri-bronchial thickening (red arrow), and evolution of the small area of consolidation to a slightly larger but less dense GGO appearance (left middle lobe, green arrow). PET/CT on the same day showed normalization of FDG-avidity in the lung infiltrates (not shown).

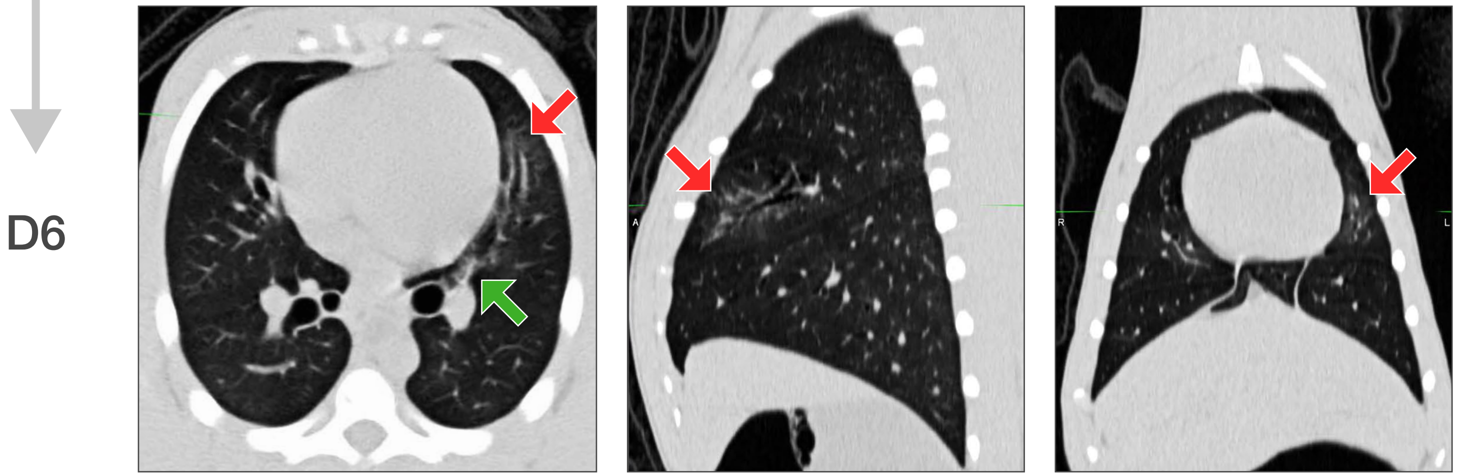


Compared to D4, chest CT on D6 showed continued improvement in the GGO (left middle lobe, red arrow) and organization and reduction in size of the GGO in the left middle lobe (green arrow).

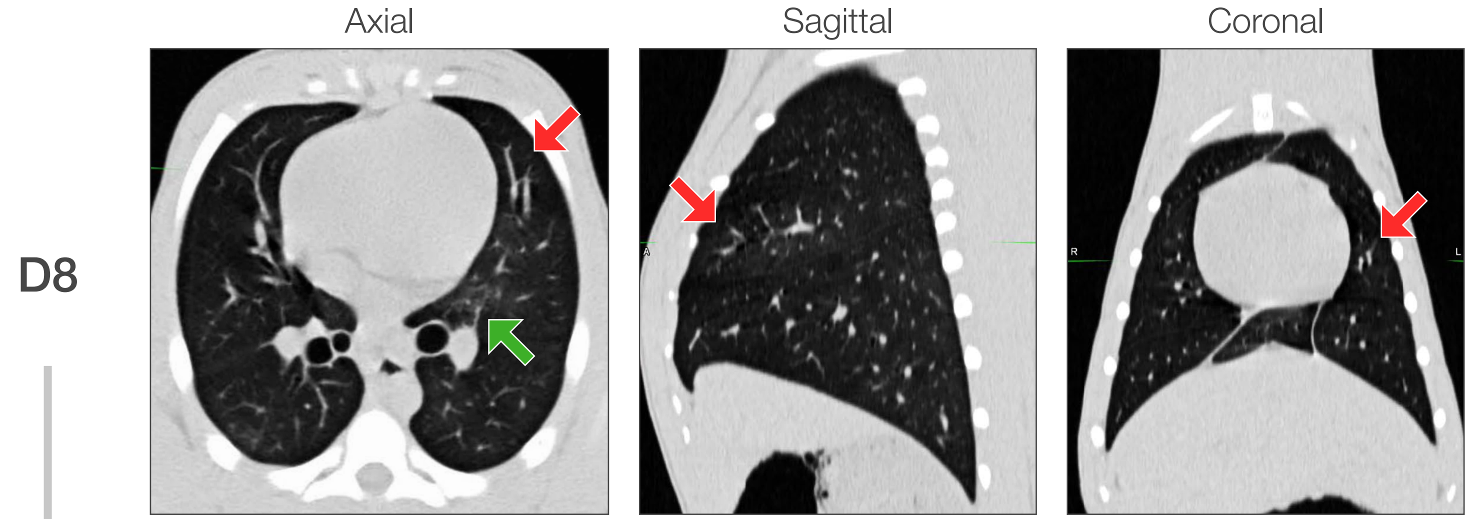


Compared to D6, chest CT on D8 showed continued improvement overall. GGO in the left middle lobe had resolved, and the central left middle lobe infiltrate was slightly larger but less dense overall, now with GGO appearance.

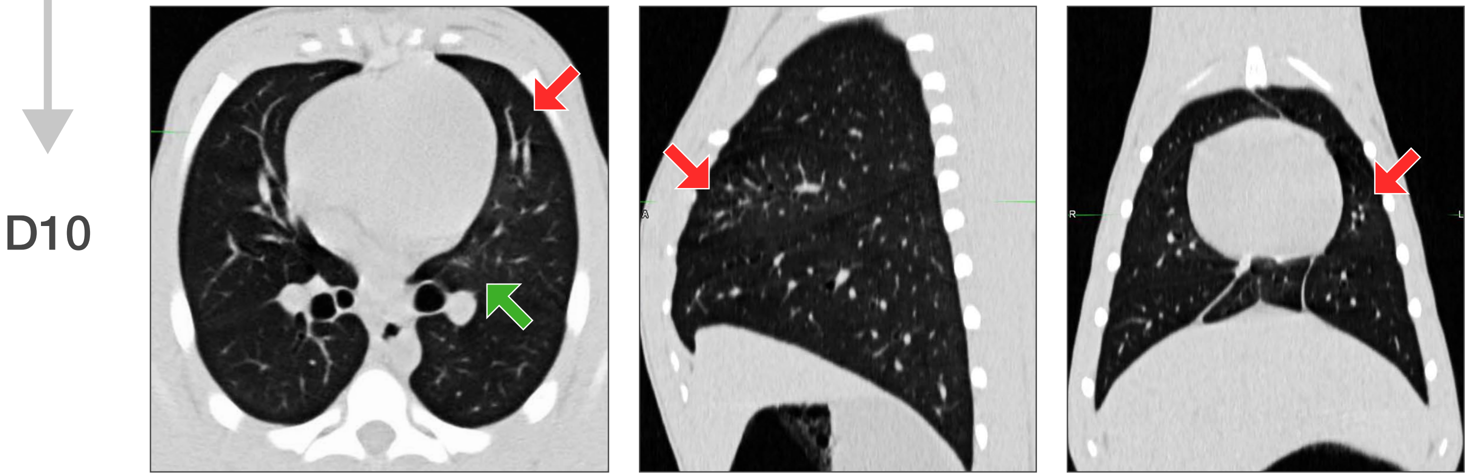


Compared to D8, chest CT on D10 showed near or full resolution of previously described abnormalities.

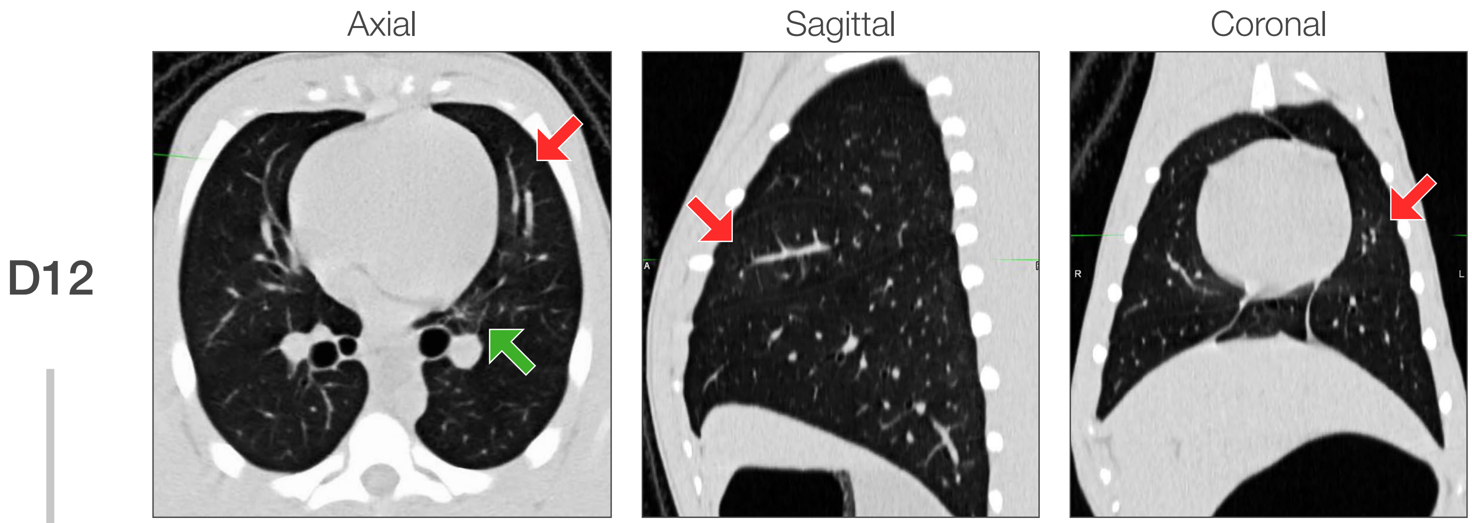


Compared to D10, chest CT on D12 showed new atelectasis in the right accessory and left middle lobe (not shown). Slight increase in the central left middle lobe GGO (green arrow) was noted.

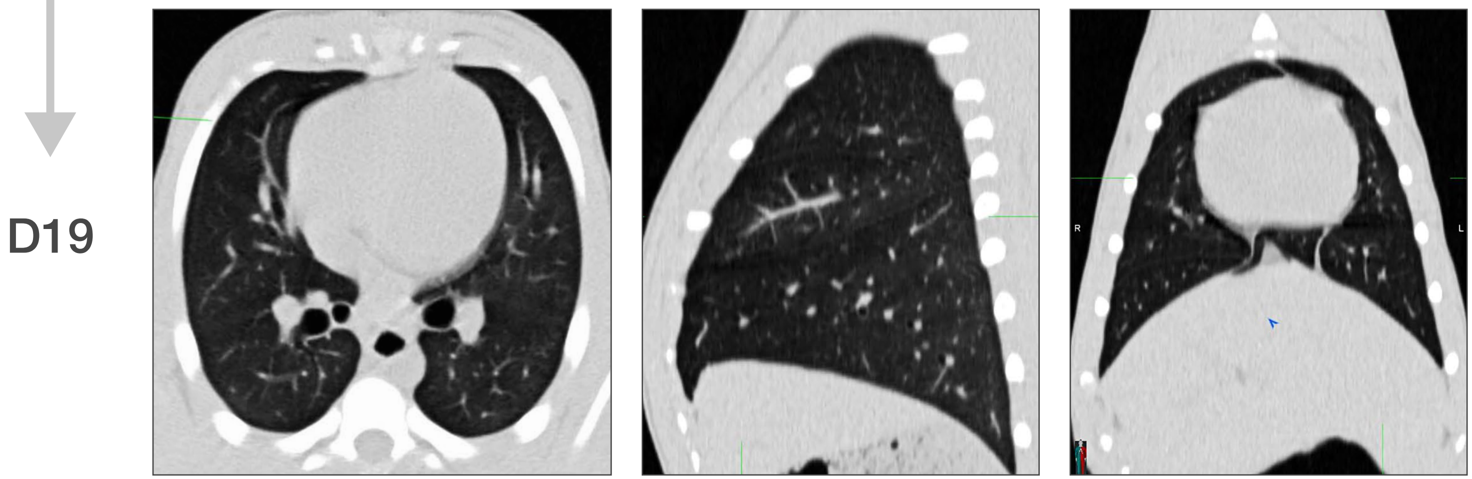


Compared to D12, chest CT on D19 showed near to full resolution of all previously noted abnormalities. Two small areas of GGO, likely atelectasis, were noted in each posterior lower lobe (not shown).

### Macaque V2

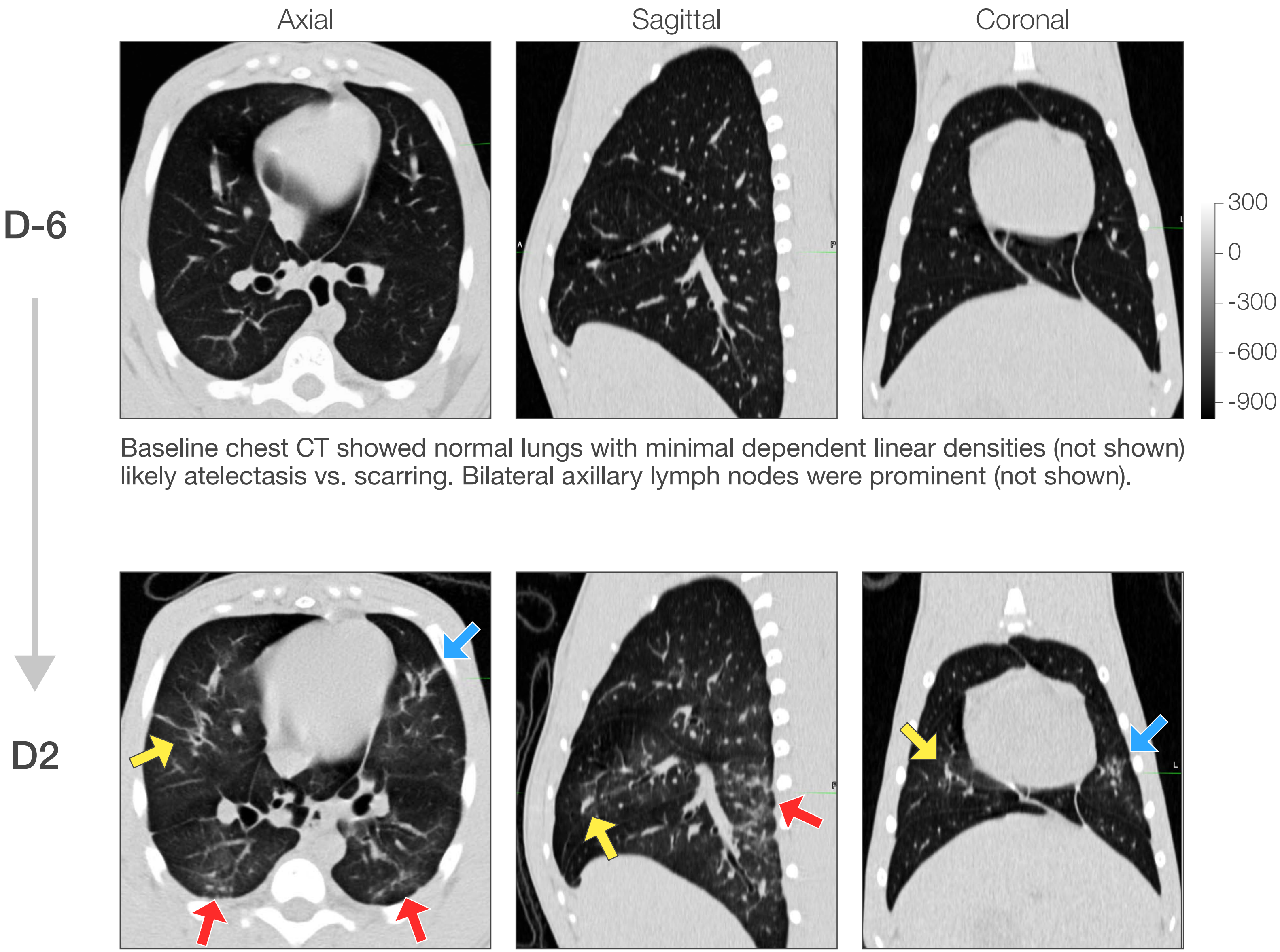


Baseline chest CT showed normal lungs with minimal dependent linear densities (not shown) likely atelectasis vs. scarring. Bilateral axillary lymph nodes were prominent (not shown).

Compared to baseline, chest CT on D2 showed new, bilateral, multi-lobar abnormalities that included: multifocal GGOs mixed with consolidation and reticulation (anterior left middle lobe, blue arrow); posterior GGO with reticulation (right lower lobe, left lower lobe, red arrows); linear opacities and bronchial wall-thickening (right middle lobe, yellow arrow) and peri-bronchial consolidation (right accessory lobe, not shown). Scattered areas of patchy GGO and linear opacity were otherwise noted (not shown). The right accessory, left lower, and multiple posterior dependent infiltrates were FDG-avid by same day PET/CT scan (not shown).

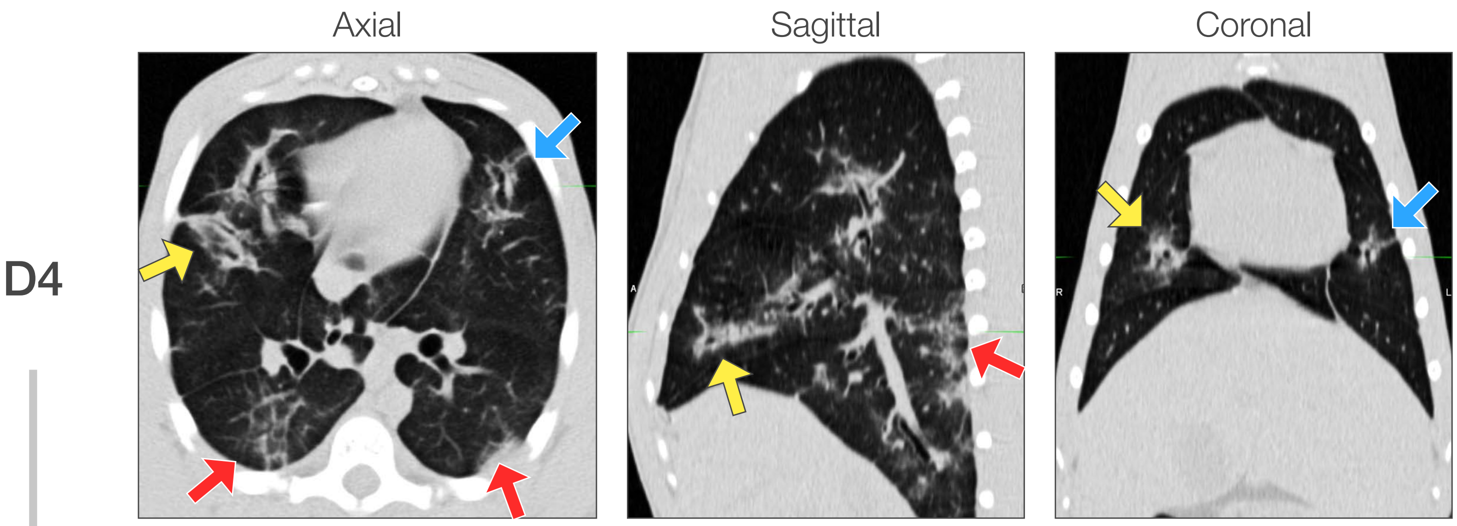


Compared to D2, chest CT on D4 showed variable progression, improvement, or a mixed response in the previously identified areas and new abnormalities. In the right lung, middle lobe peri-bronchial thickening and consolidation (yellow arrow) had progressed; the right accessory lobe abnormalities had variably expanding or coalescing infiltrates (not shown); there was new central peri-bronchial GGO in the right upper lobe (not shown). In the left lung, the left middle lobe showed subtly increased GGO (blue arrow) while the central lower lobe infiltrate was stable. New intra-septal thickening ("crazy-paving") on top of increased GGO developed in the posterior right lung (red arrow), with organization and subpleural thickening noted on the left (red arrow).

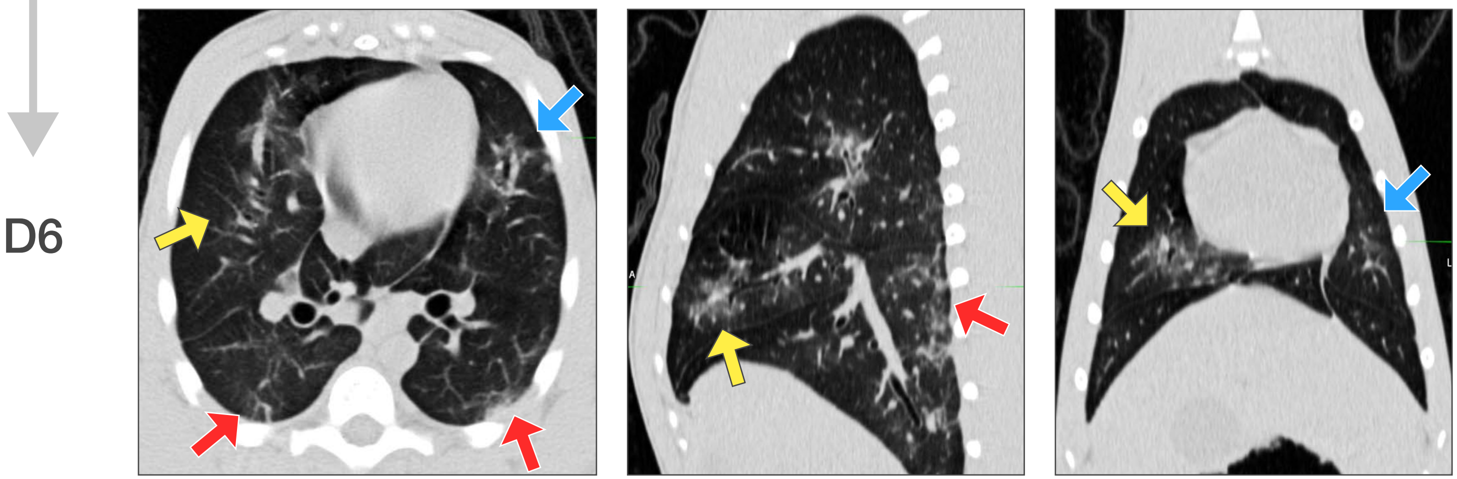


Compared to D4, chest CT on D6 showed persistence or improvement in some abnormalities, including in the right middle lobe (yellow arrow), the bilateral posterior changes (red arrows), and in the left middle lobe (blue arrow). There were several new pleural-based lesions in both upper lobes (not shown). PET/CT from the same day (not shown, refer to appropriate figure) showed increased FDG-avidity in many of these lung abnormalities compared to D2 PET/CT.

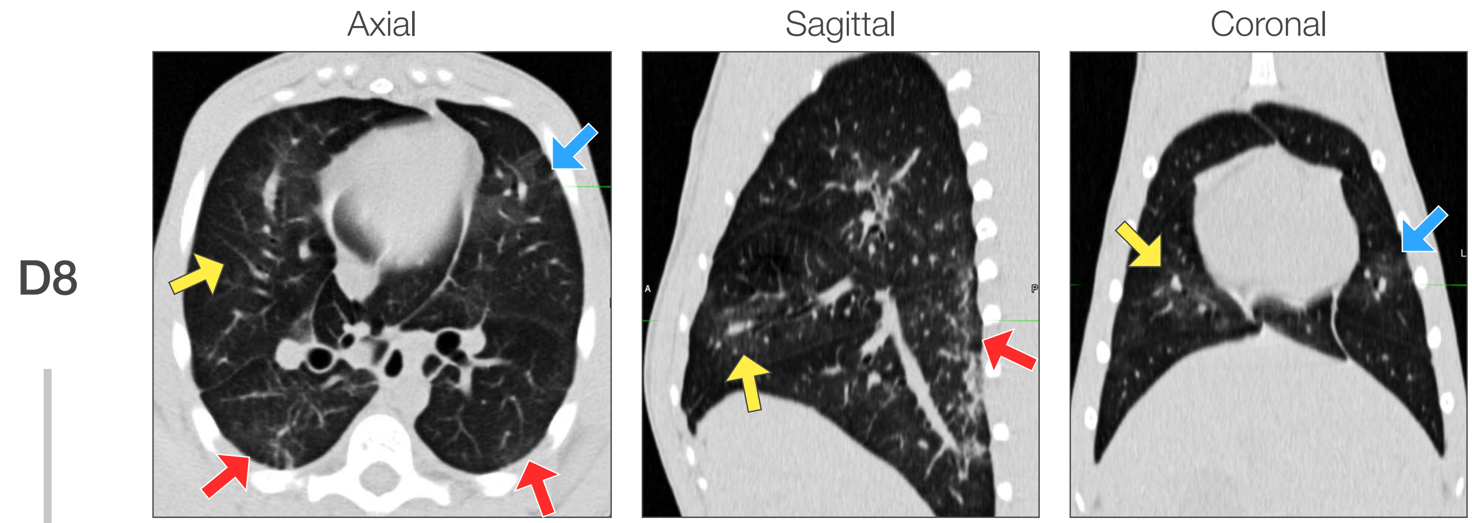


Compared to D6, chest CT on D8 showed general improvement or resolution in noted abnormalities and no progression. The bilateral posterior GGO or reticular abnormalities (red arrows) were improved or unchanged.

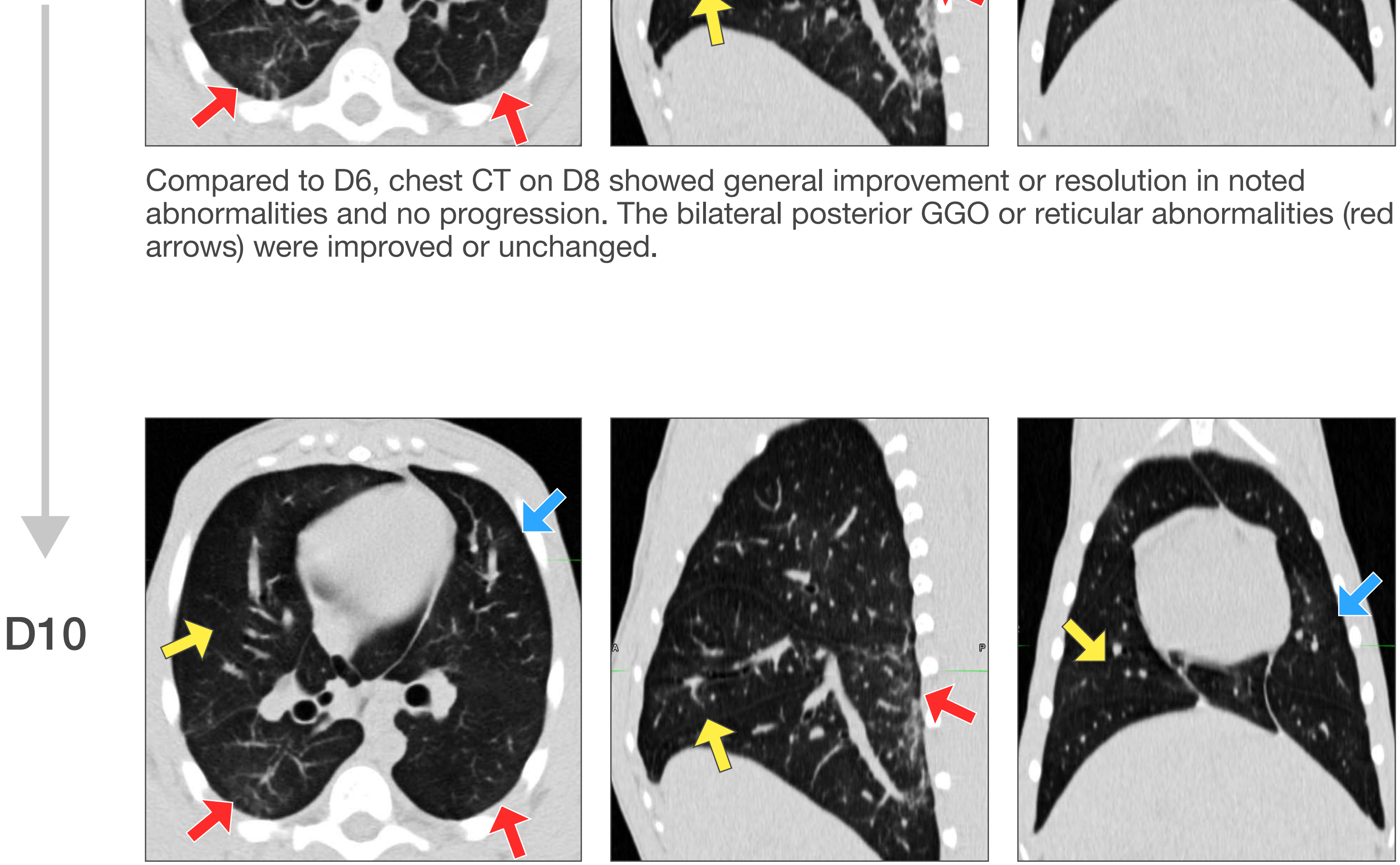


Compared to D8, chest CT on D10 showed continued improvement or resolution of noted abnormalities. The right upper lobe central GGO had resolved; minimal residual GGO remained in the right middle lobe (yellow arrow) and bilateral upper lobe pleural based abnormalities (not shown); posterior reticulations continued to improve but persisted (red arrow, sagittal).

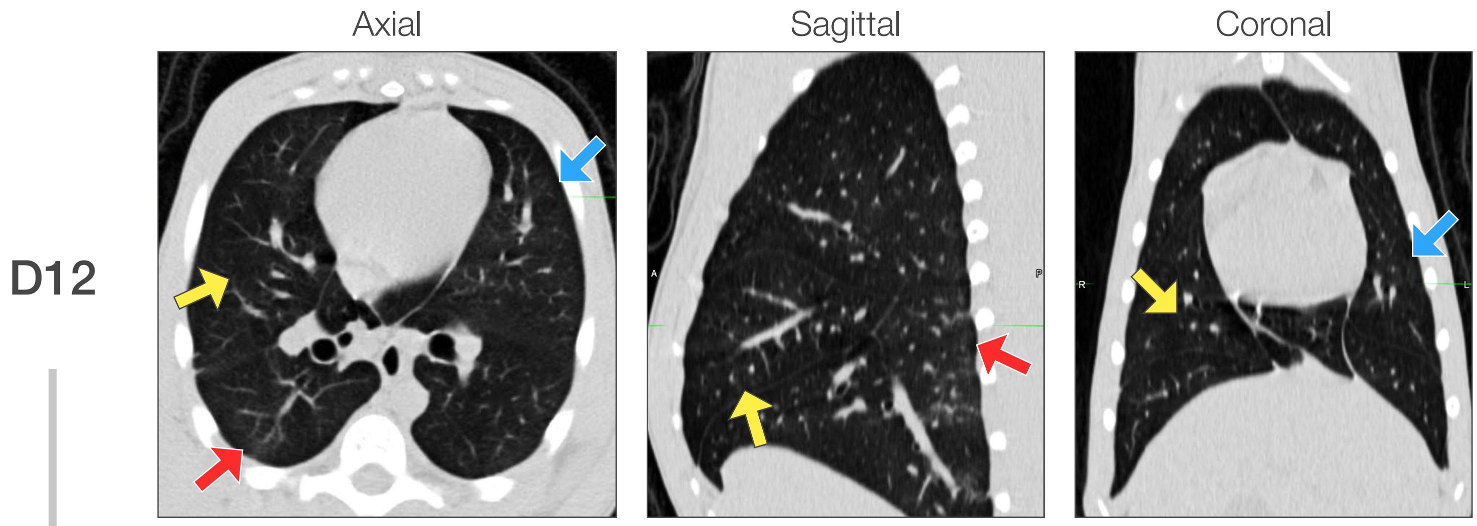


Compared to D10, chest CT on D12 showed improvement to near-resolution in all noted abnormalities without progression or new changes. Though improved, mild to minimal posterior reticulations persisted.

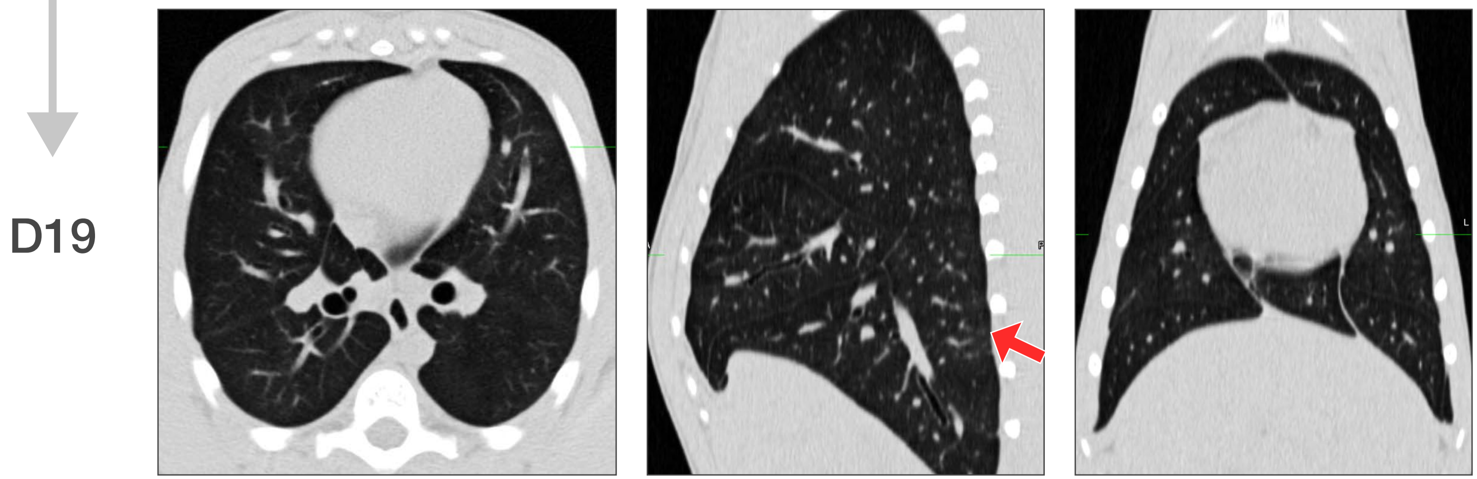


Compared to D12, chest CT on D19 showed continued improvement to near-resolution, with minimal residual posterior right lower lobe reticulations (red arrow) but no progression or new abnormalities.

### Macaque V3

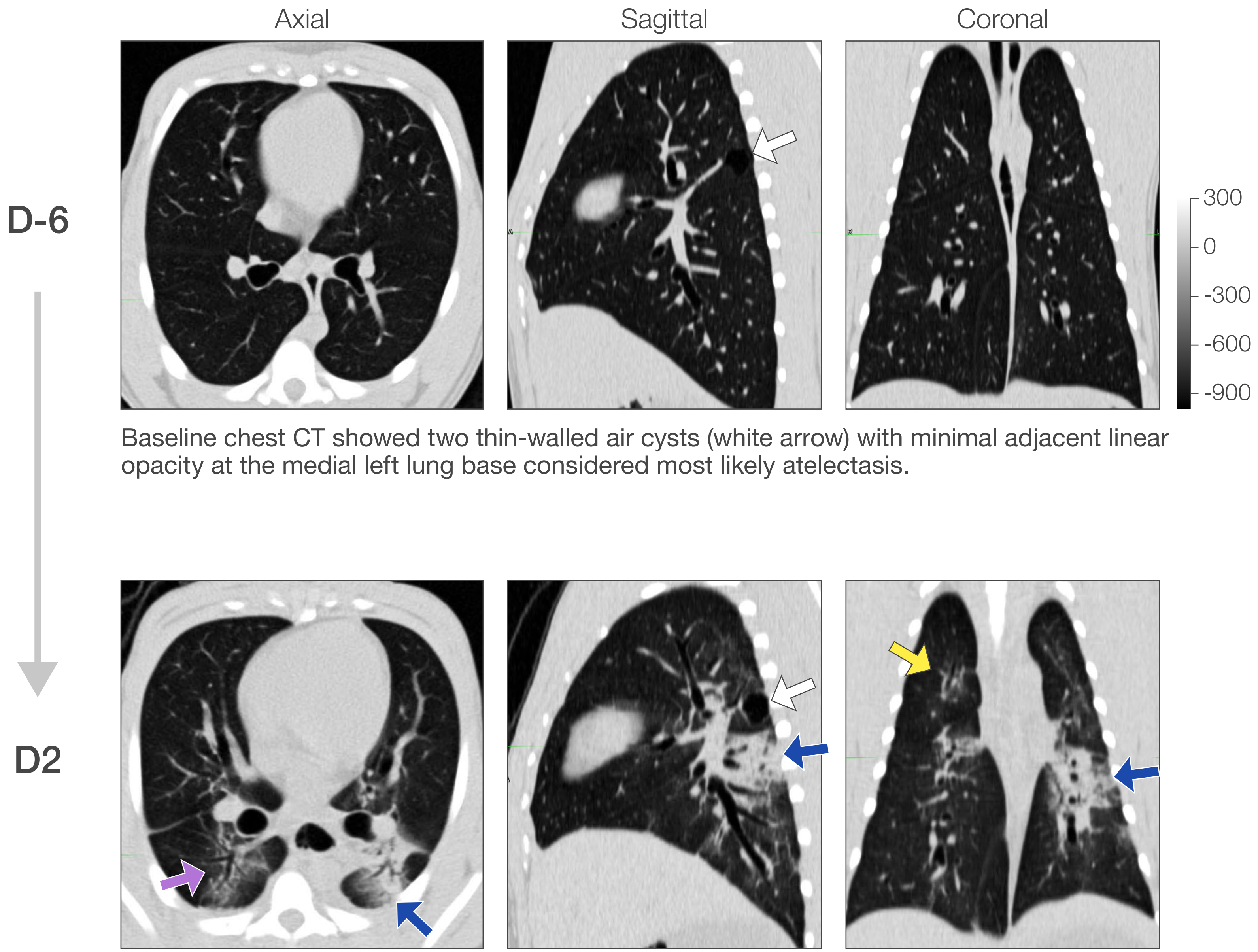


Baseline chest CT showed two thin-walled air cysts (white arrow) with minimal adjacent linear opacity at the medial left lung base considered most likely atelectasis.

Compared to baseline, chest CT on D2 after inoculation showed new, bilateral, multi-lobar abnormalities. In the right lung, right upper lobe GGO with superimposed inter-septal thickening or "crazy paving" (yellow arrow), right accessory lobe GGO (not shown), and right lower lobe central peri-bronchial infiltrate (purple arrow) and sub-pleural band (not shown) were noted. In the left lung, dense airspace consolidation with multiple air bronchograms was noted in both the central and peripheral left lower lobe (blue arrow). These structural abnormalities (infiltrates) were FDG-avid by PET/CT on the same day.

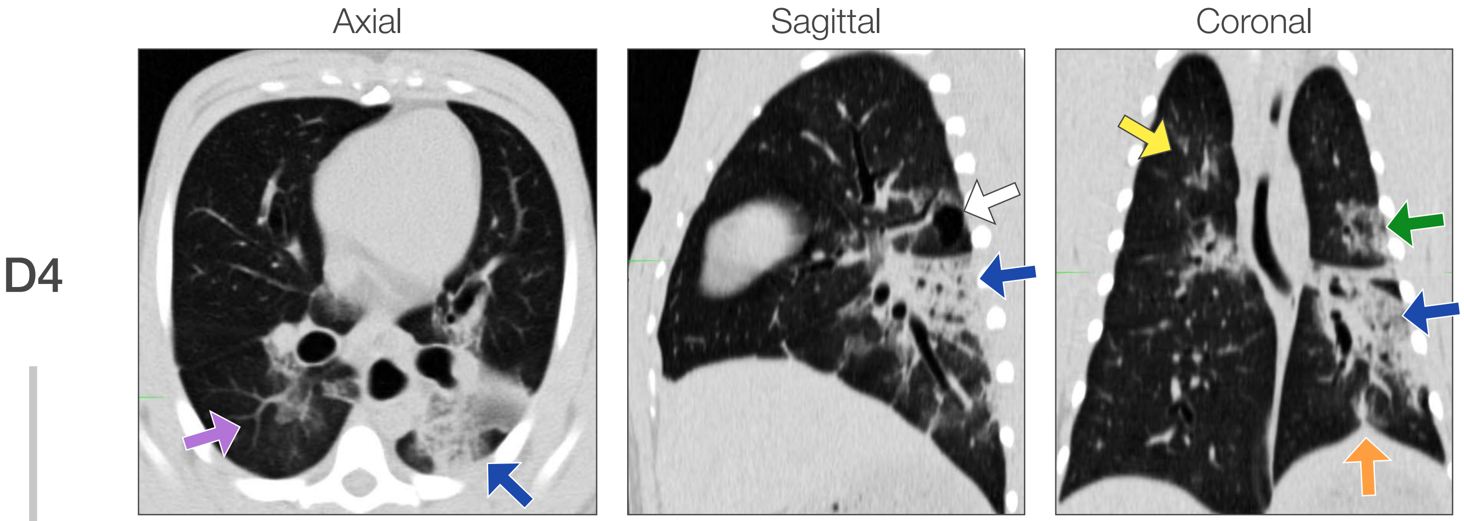


Compared to D2, chest CT on D4 showed progression of bilateral multi-lobar infiltrates, most notably with peri-bronchial, peripheral, and caudal extension of the left lower lobe consolidation (blue arrow), and extension of right upper lobe peri-bronchial consolidation and crazy-paving (yellow arrow). New peripheral infiltrate was noted subpleurally in the medial posterior left lower lobe (orange arrow), the inferior-posterior left upper lobe (green arrow), and new GGO was noted in the central left upper lobe and the right accessory lobe. Mixed evolution of the infiltrate in the superior right lower lobe was noted with organizing consolidation medially and improvement centrally in GGO and paving (purple arrow).

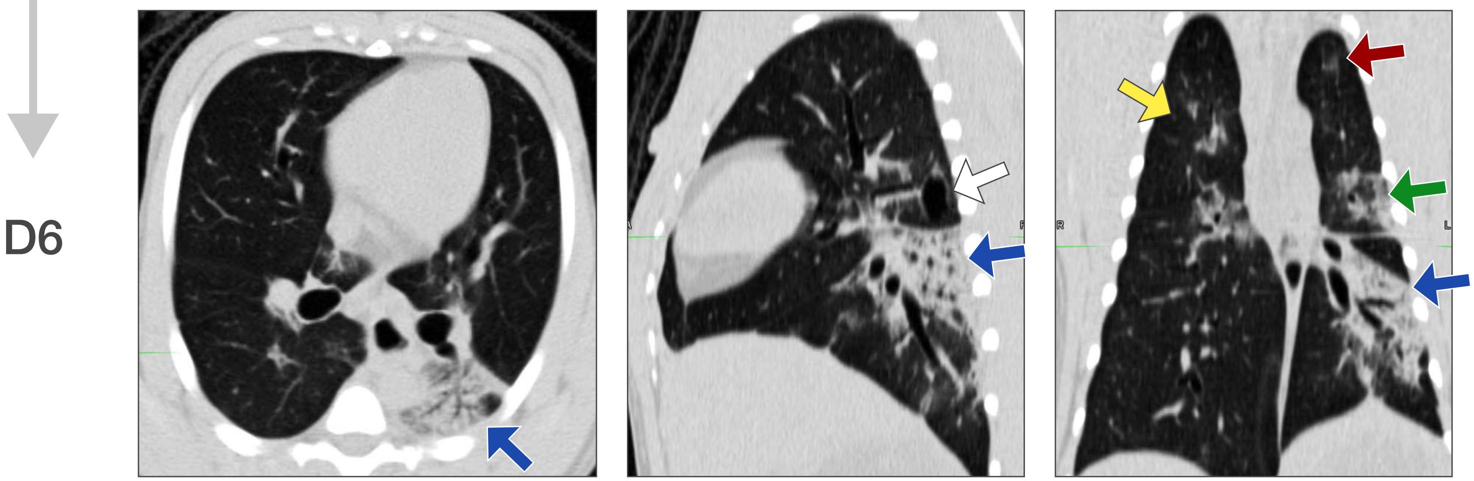


Compared to D4, chest CT on D6 showed persistence of the previously noted structural abnormalities, with tendency to a slight overall increase in consolidation (blue arrow, green arrow). The left upper lobe GGO was increased (maroon arrow). Notably, PET/CT done on the same day (not shown, see figure 4) showed a marked increase in FDG-avidity in these abnormal areas when compared to D2 PET/CT, particularly in the left lower lobe (blue arrow) and the right upper lobe.

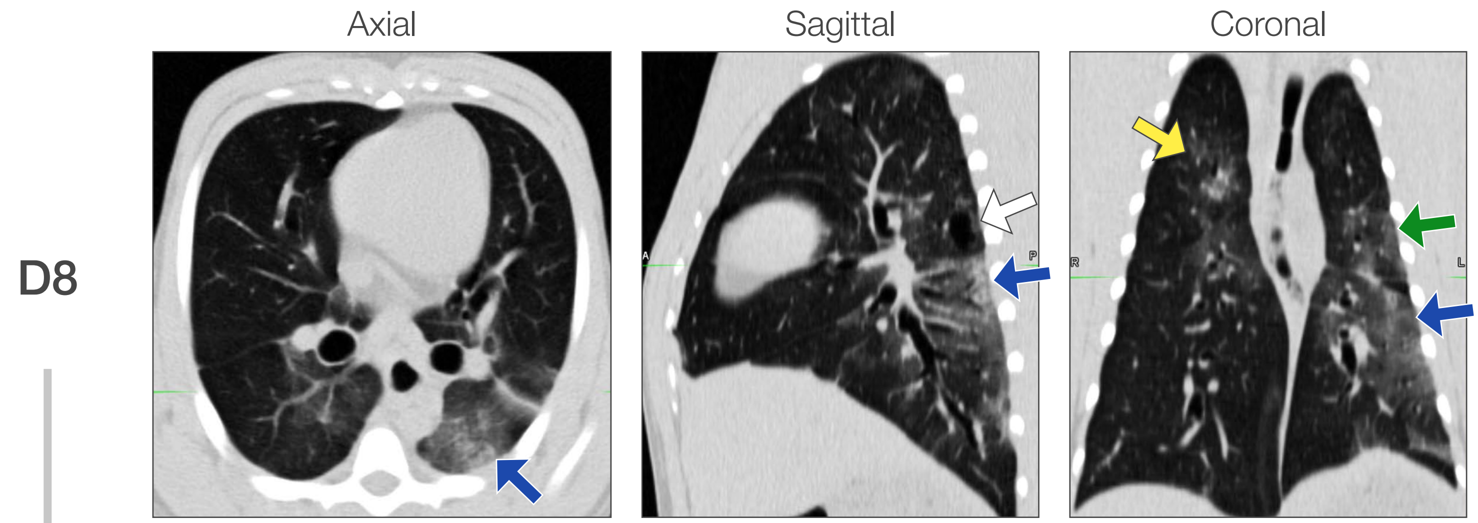


Compared to D6, chest CT on D8 showed evolution of almost all abnormalities from more dense consolidation to less dense GGO. This was most notable in the left lower lobe (blue arrow) which increased in overall size but was much less dense. It was also apparent in the inferior posterior left upper lobe (green arrow) and right upper lobe (not clearly shown).

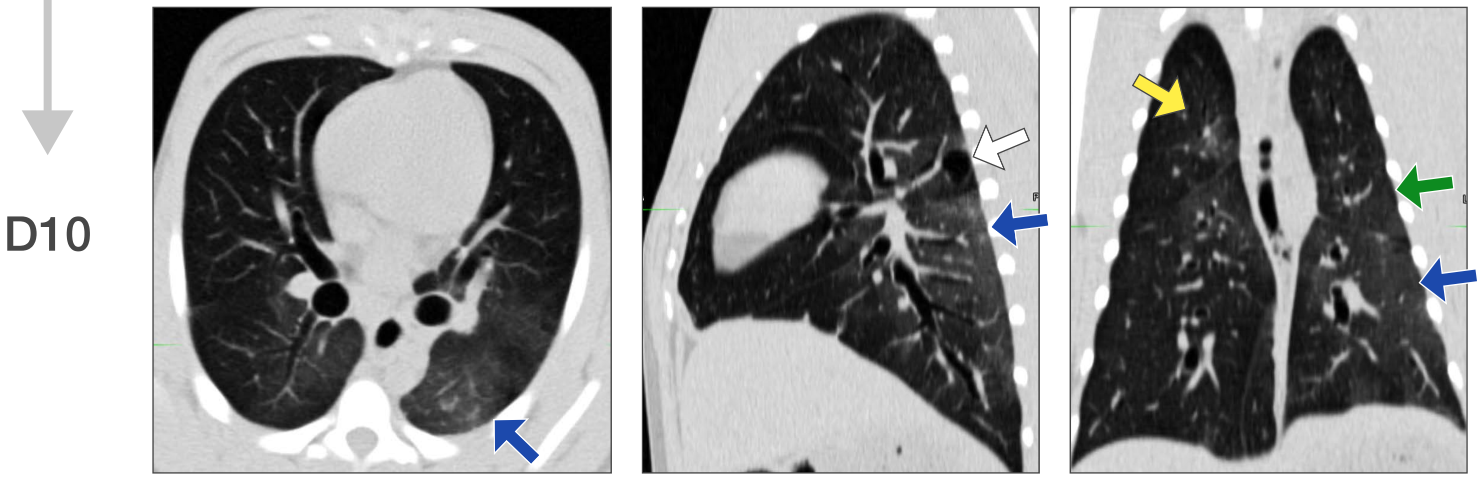


Compared to D8, chest CT on D10 showed continued improvement in all noted abnormalities, with evolution towards less dense GGO and linear opacity bands suggestive of organizing pulmonary remodeling (not shown).

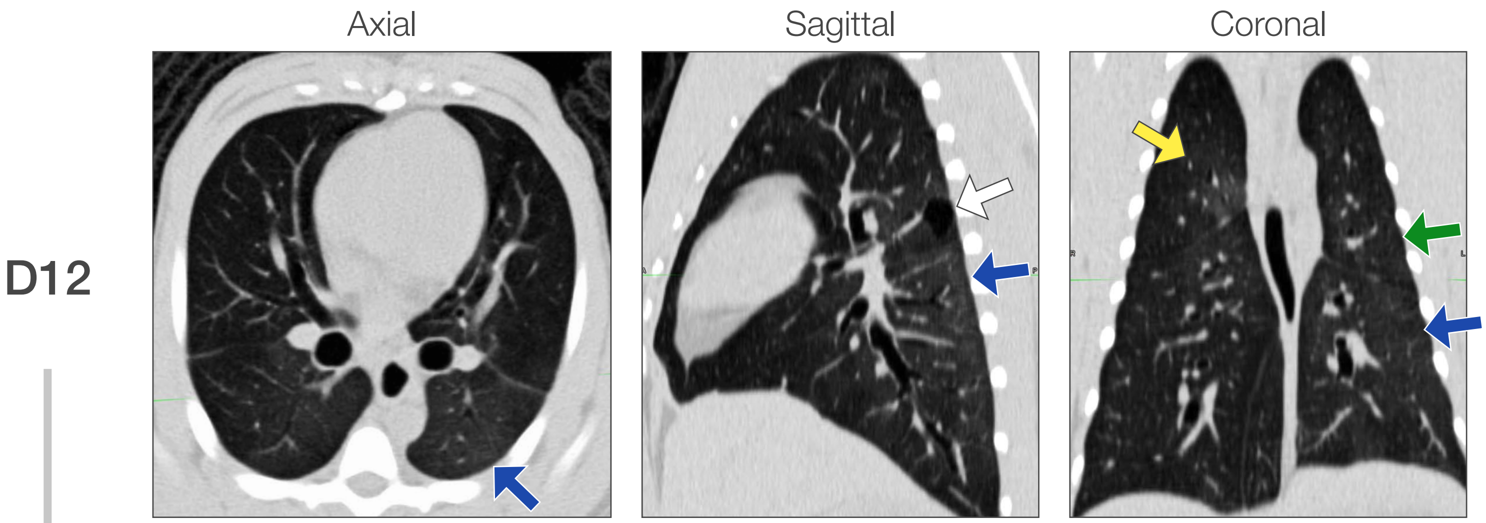


Compared to D10, chest CT on D12 showed continued decrease in radiodensity of previously noted residual GGOs and linear opacities.

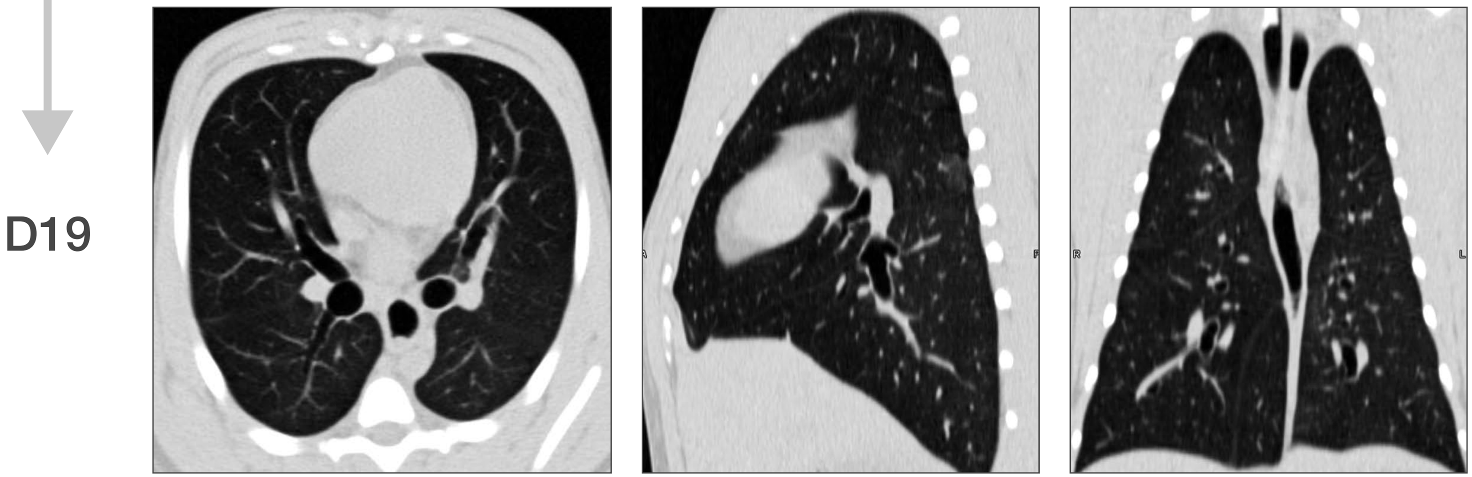


Compared to D12, chest CT on D19 showed improvement to near resolution in all previously noted abnormalities and no new abnormalities.
