## Supplemental Figure 5 for "Characteristic and quantifiable COVID-19-like abnormalities in CT- and PET/CT-imaged lungs of SARS-CoV-2-infected crab-eating macaques (*Macaca fascicularis*)"

Supplementary Figure 5

### Macaque V1

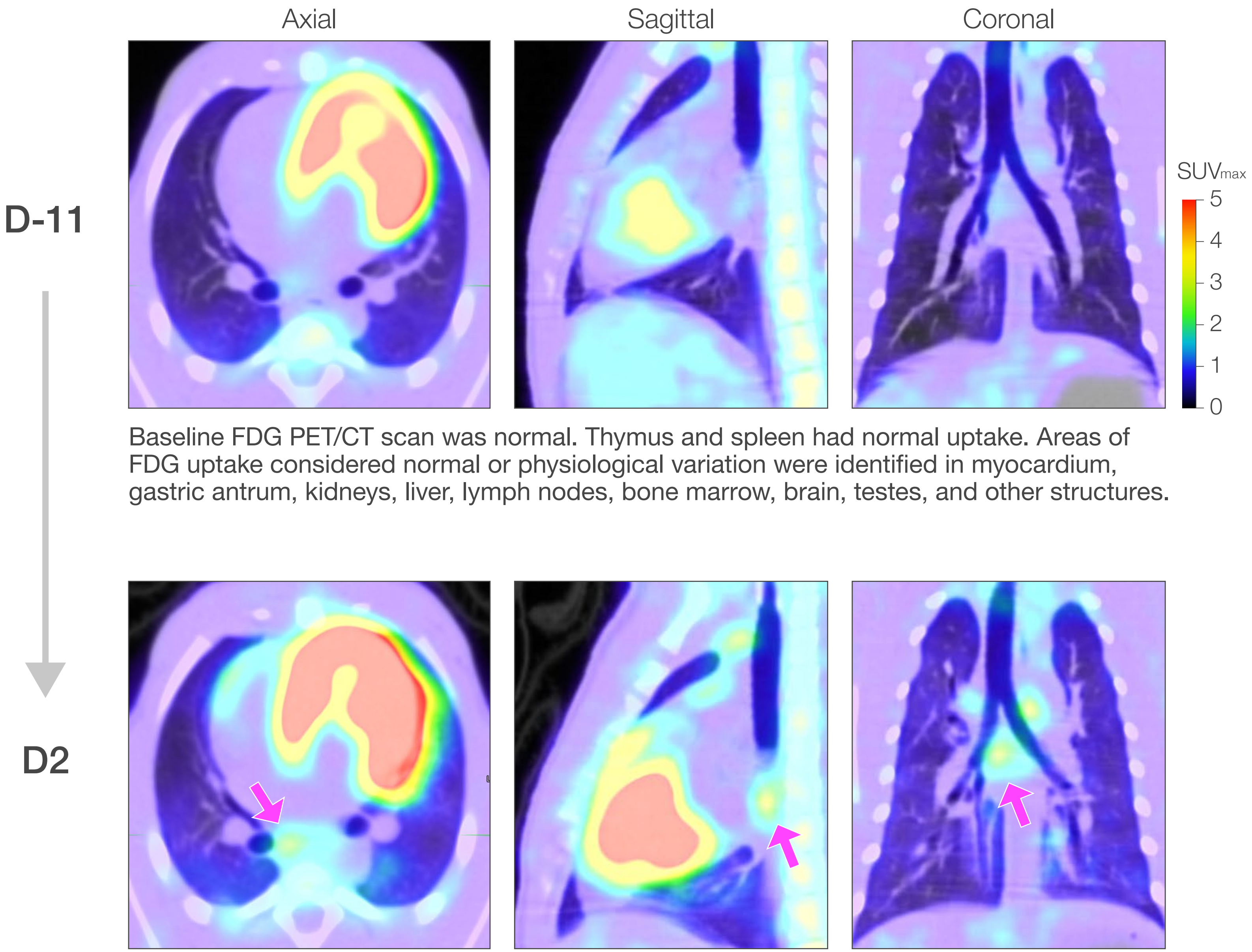


Baseline FDG PET/CT scan was normal. Thymus and spleen had normal uptake. Areas of FDG uptake considered normal or physiological variation were identified in myocardium, gastric antrum, kidneys, liver, lymph nodes, bone marrow, brain, testes, and other structures.

Compared to baseline, FDG PET/CT on D2 after inoculation showed new FDG uptake in the lung parenchyma corresponding to structural abnormalities identified on CT scan. These included the right upper lobe, left lower lobe, and other posterior and dependent areas of the lung (not shown). New FDG uptake was noted in the bilateral paratracheal and subcarinal nodes in the mediastinum (pink arrow). Increased uptake noted in bilateral axillary and supra-clavicular areas was difficult to distinguish between lymph nodes and brown fat. A subtle increase in thymus uptake was noted, and a decrease in FDG signal in the brain, liver, and spleen (not shown).

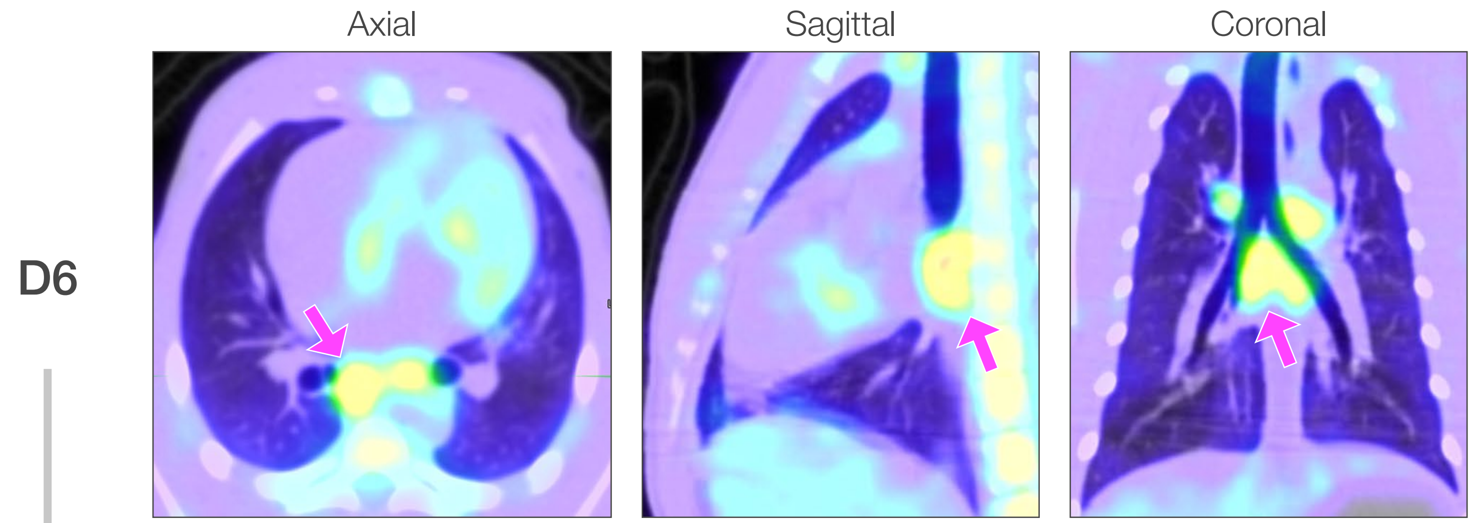


Compared to D2, FDG PET/CT on D6 after inoculation showed normalization of FDG uptake in the areas associated with CT lung abnormalities, with particular improvement in the left lung (not shown). There was marked increase in FDG uptake in intra-thoracic hilar and mediastinal lymph nodes (pink arrow). A less marked increase in FDG uptake was seen in the spleen, brain, and liver, while decreased uptake was noted in the myocardium and other regions of the brain. Persistent but reduced uptake in brown fat regions was noted.

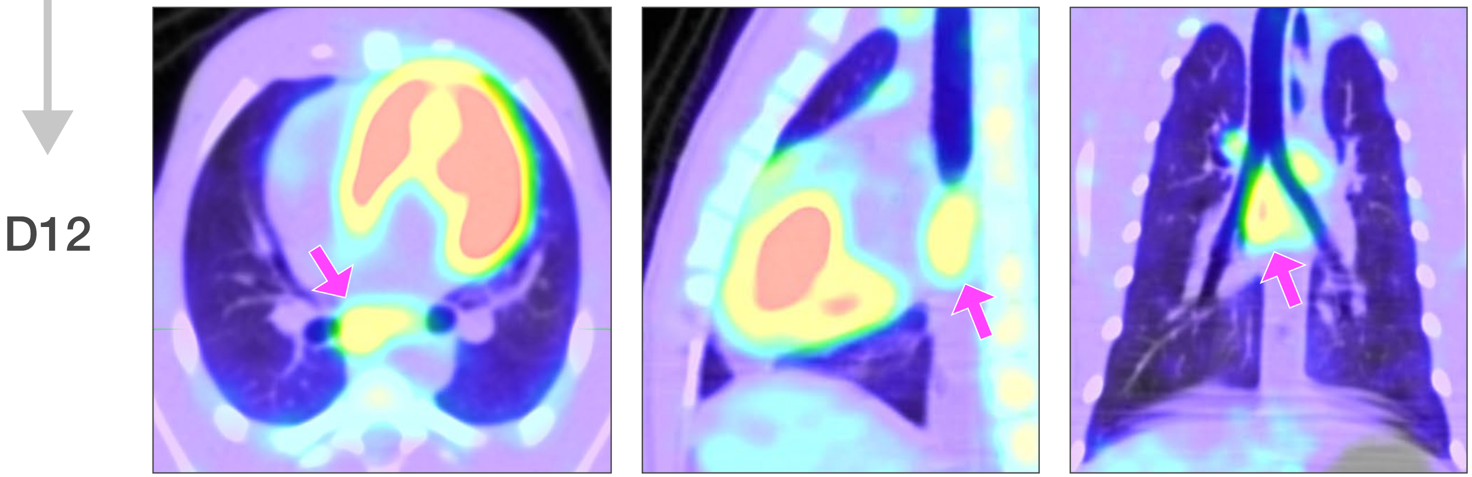


Compared to D6, FDG PET/CT on D12 showed no significant lung uptake, a variable increase in uptake in subcarinal lymph nodes (increased on right, decreased on left) and decreased uptake in the left parabronchial lymph node. A decrease in FDG uptake was also noted in the spleen, thymus (slight), spinal bone marrow, and brain.

### Macaque V2

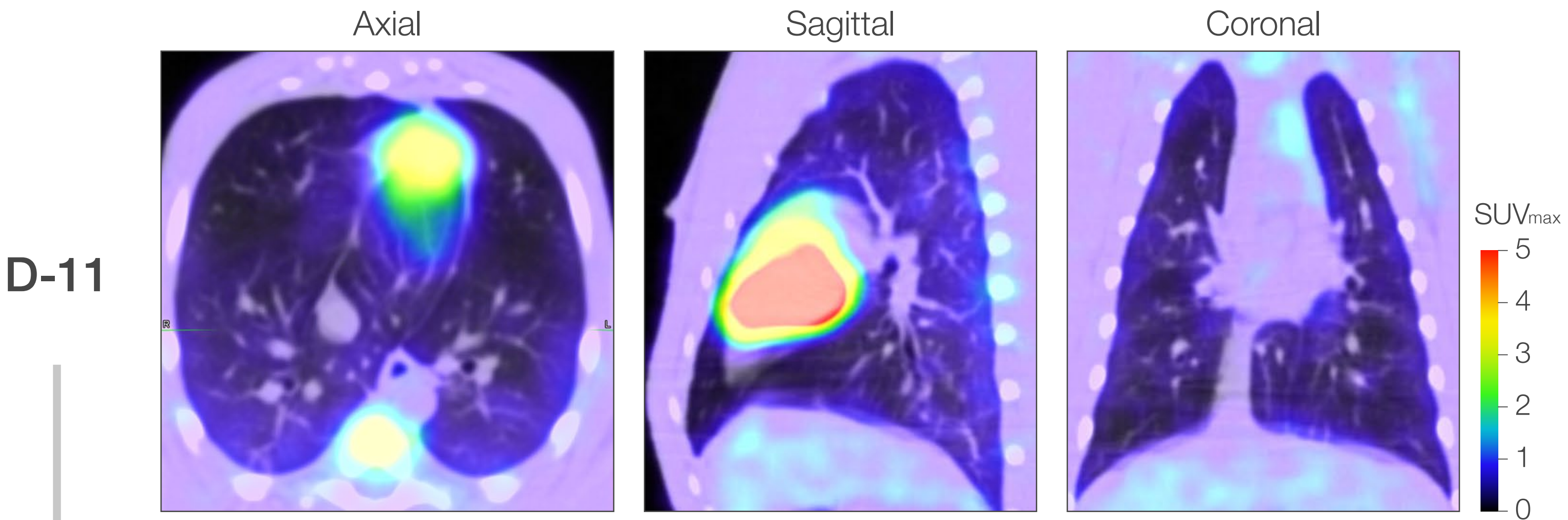


Baseline FDG PET/CT scan showed prominent but likely normal thymus and spleen uptake. Bone marrow uptake in the pelvis was more than typically seen. Areas of FDG uptake considered normal or physiological variation were identified in myocardium, gastric antrum, kidneys, liver, lymph nodes, bone marrow, brain, testes, and other structures.

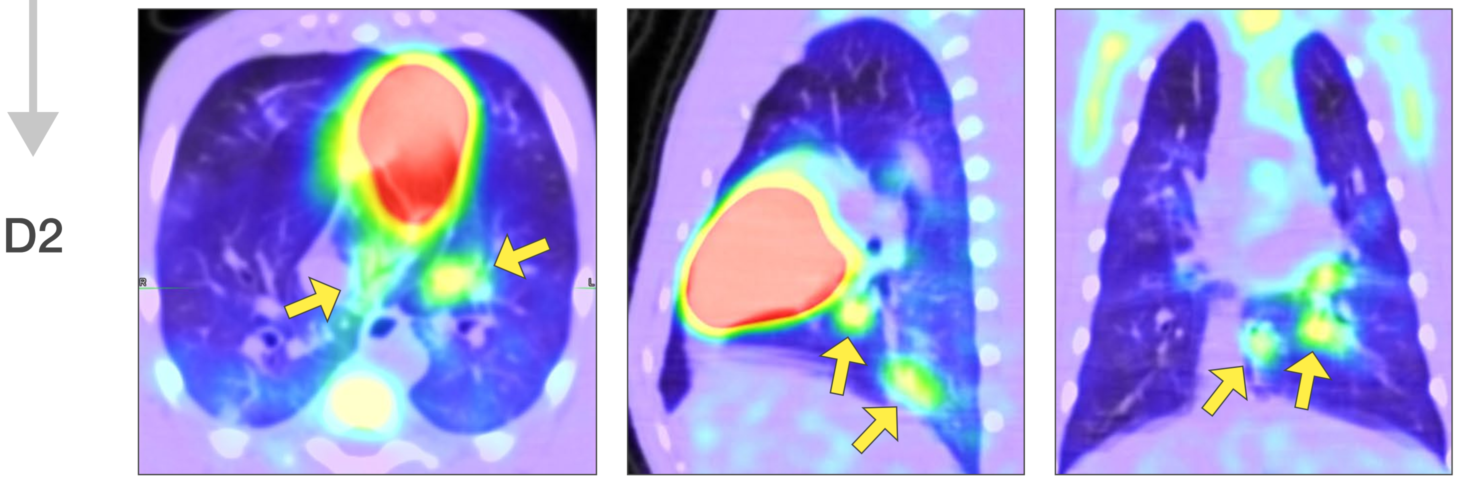


Compared to baseline, FDG PET/CT on D2 after inoculation showed new FDG uptake in the lung parenchyma corresponding to structural abnormalities identified on CT scan. FDG-avidity was noted in new lung infiltrates (yellow arrows) in the right accessory lobe, left lower lobe and multiple other smaller areas in the posterior and dependent lungs. New FDG uptake was noted in the bilateral paratracheal, hilar, and subcarinal lymph nodes. Increased uptake was noted in bilateral axillary, neck, and supraclavicular areas thought likely a combination of lymph nodes and brown fat. A decrease in FDG uptake was noted in thymus, (subtle), brain, liver, and spleen.

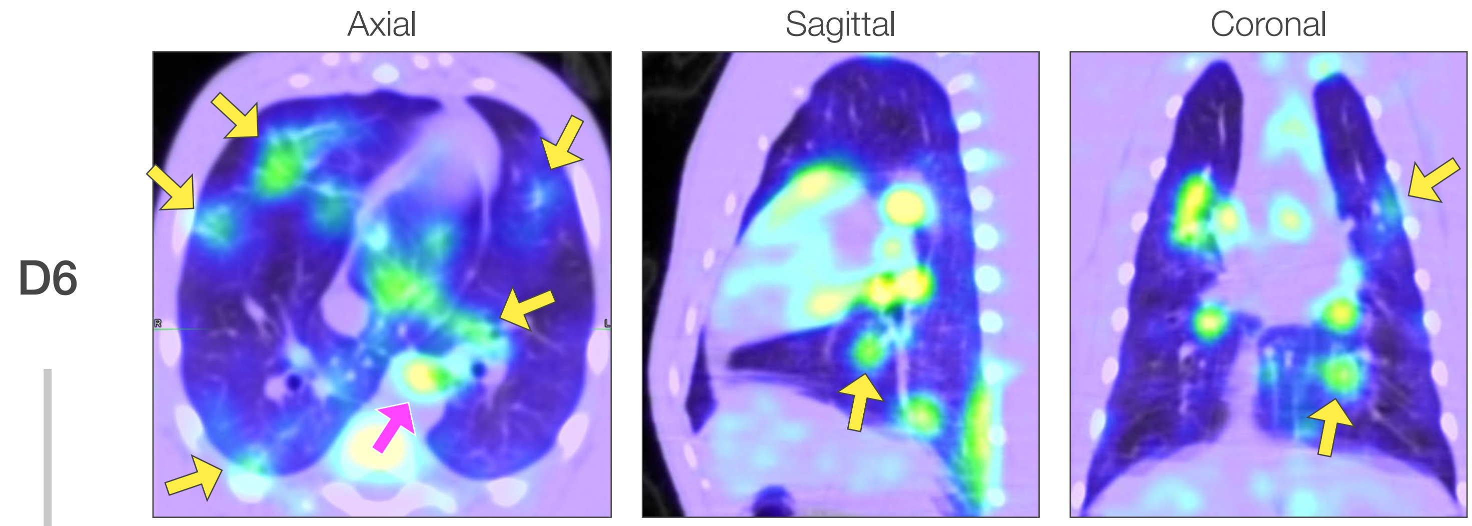


Compared to D2, FDG PET/CT on D6 after inoculation showed a marked increase in FDG uptake in intrathoracic lymph nodes (pink arrow) as well increased uptake in multiple previously described (right accessory lobe, left lower lobe) and new lung abnormalities (peripheral upper lobes; yellow arrows). An increase in FDG uptake was also noted in thymus, spleen, bone marrow, brain, and liver with a decrease in myocardium and brown fat regions. FDG uptake in other organs was thought to be normal or physiologic variation.

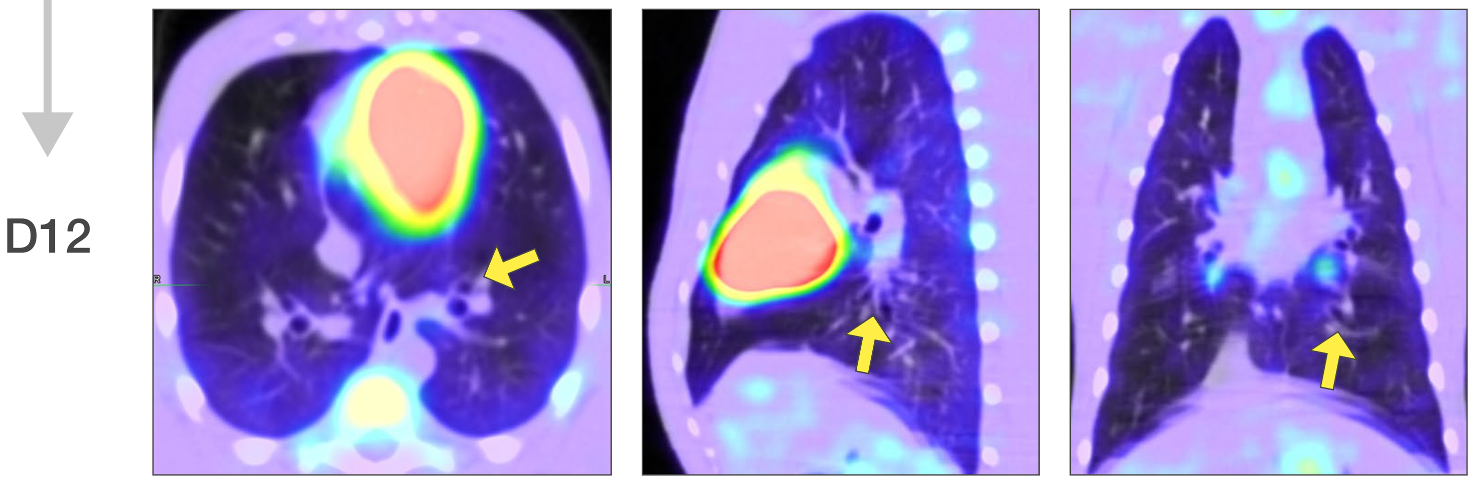


Compared to D6, FDG PET/CT on D12 after inoculation showed resolution or decrease in all FDG-avid lesions, including all lung and mediastinal and hilar lymph nodes. The highest residual signal was in the right subcarinal lymph node. FDG uptake was decreased in the spleen and considered normal or of physiologic variation in other organs.

### Macaque V3

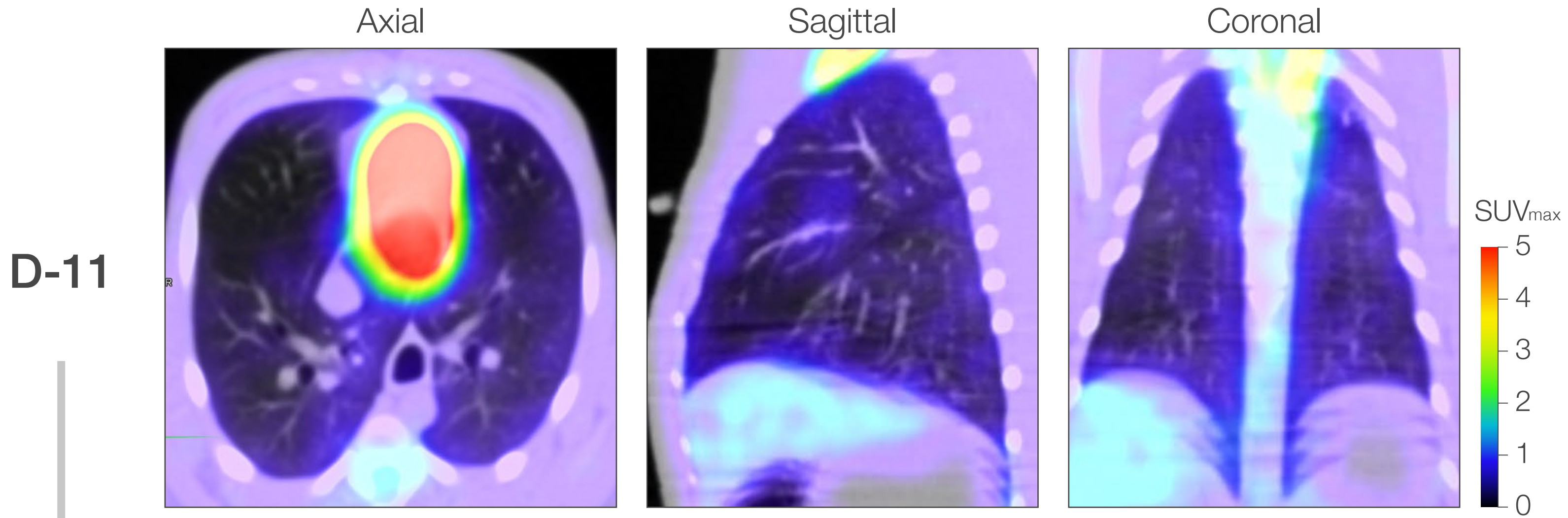


Baseline FDG PET/CT scan 11 days prior to inoculation showed massive increase in FDG uptake in the neck and upper thorax, predominantly brown fat, though indistinguishable from adjacent lymph nodes or other structures. Thymus uptake could not be distinguished from brown fat uptake. The spleen had low normal uptake. Areas of FDG uptake considered normal or physiological variation were identified in myocardium, gastric antrum, kidneys, liver, lymph nodes, bone marrow, brain, testes, and other structures.

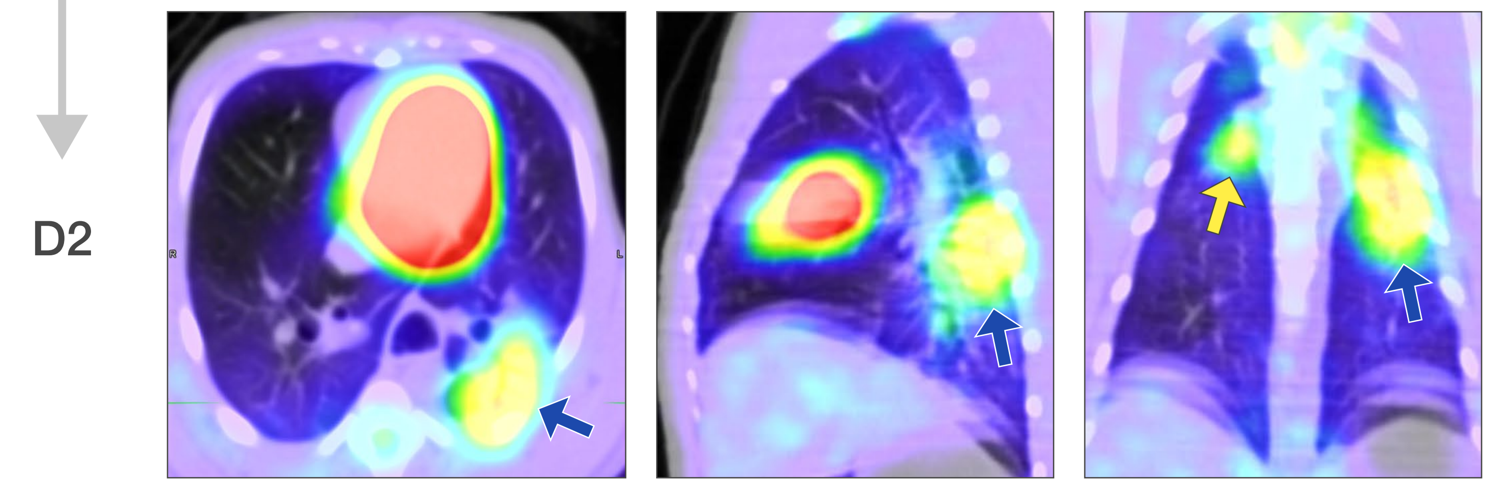


Compared to baseline, FDG PET/CT on D2 after inoculation showed new FDG uptake in the lung parenchyma corresponding to structural abnormalities identified on CT scan. FDG-avidity was noted in multiple new lung infiltrates including densely consolidated areas of the left lower lobe (blue arrow), posterior infiltrate in the right lower lobe (not shown), and right upper lobes (yellow arrow). New mediastinal FDG uptake was noted in the bilateral hilar and subcarinal lymph nodes. There was decreased but still substantial residual brown fat activation in the upper thorax. An increase in FDG uptake was noted in bone marrow, spleen (subtle), and thymus (though difficult to distinguish from myocardium and brown fat signal). A decrease was noted in brain and liver.

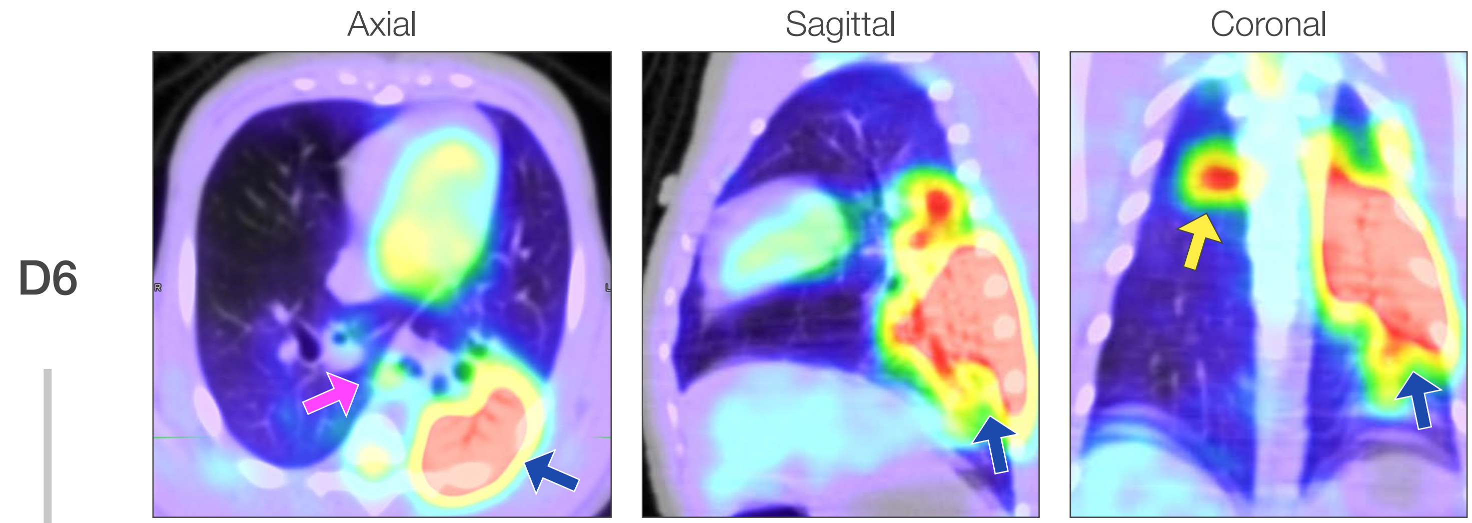


Compared to day 2, 18-FDG PET/CT on day 6 after inoculation showed a marked increase in FDG uptake in previously described lung abnormalities in the left lower lobe (blue arrow), right lower lobe (yellow arrow) right upper lobe (not shown) as well as in intra-thoracic bilateral hilar and mediastinal lymph nodes (pink arrow). A less marked increased in uptake was noted in spleen, bone marrow, brain, and liver. A decrease was noted in myocardium and thymus, and mixed changes in persistent areas of brown fat activation. FDG uptake in other organs was thought to be normal or physiologic variation.

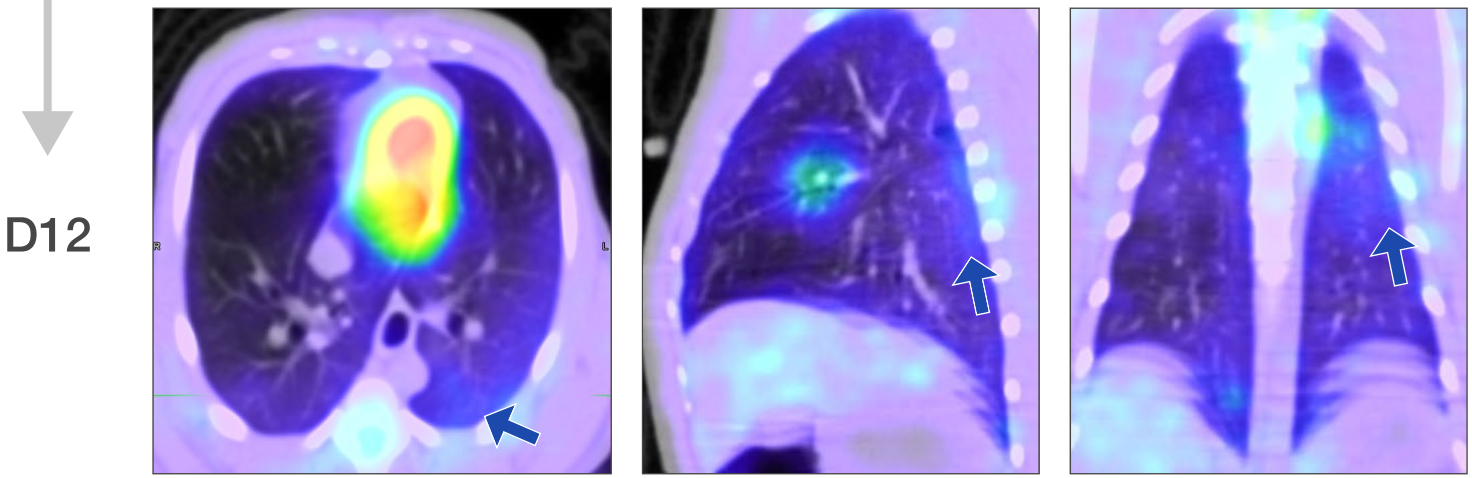


Compared to D6, FDG PET/CT on D12 after inoculation showed resolution or decrease in FDG uptake in all lung areas and mediastinal and hilar lymph nodes. The highest residual signal was in the right subcarinal lymph node. FDG uptake was decreased in the bone marrow, thymus and spleen, and considered normal or of physiologic variation.
