## Supplemental Table 1 for "Characteristic and quantifiable COVID-19-like abnormalities in CT- and PET/CT-imaged lungs of SARS-CoV-2-infected crab-eating macaques (*Macaca fascicularis*)"

**Supplementary Table 1 | Crab-eating macaque (*Macaca fascicularis* Raffles, 1821) information**

| **Group** | **Macaque ID** | **Age** | **Weight at baseline (kg)** | **Sex** | **Inoculum** |
|---|---|---|---|---|---|
| Mock (M) | M1 | 4 years 4 months | 3.29 | F | DMEM + 2% heat-inactivated FBS |
| | M2 | 3 years 11 months | 3.17 | F | |
| | M3 | 3 years 9 months | 4.87 | M | |
| Virus (V) | V1 | 4 years 3 months | 4.34 | M | DMEM + 2% heat-inactivated FBS + $3.65 \times 10^6$ pfu SARS-CoV-2 |
| | V2 | 4 years | 4.62 | M | |
| | V3 | 4 years 4 months | 3.17 | F | |
