## Supplemental Table 4 for "Characteristic and quantifiable COVID-19-like abnormalities in CT- and PET/CT-imaged lungs of SARS-CoV-2-infected crab-eating macaques (*Macaca fascicularis*)"

**Clinical Scoring**

| Cage-Side Assessment | | |
|---|---|---|
| **Parameter** | **Description** | **Score** |
| Responsiveness and recumbency | Normal—bright, alert, responsive | 0 |
| | Mild—slightly depressed, acts disinterested when personnel in room, lying down in cage but gets up when approached | 2 |
| | Moderate/obtunded—non-responsive, very disinterested in personnel, hunched or lying down, will get up when prodded, pinched, or similarly stimulated | 4 |
| | Severe/comatose—lying down completely unresponsive to stimuli | 6 |
| Discharges | Nasal or ocular | 2 |
| | Nasal and ocular | 4 |
| Skin | Rash | 1 |
| Respiration, dyspnea, and cough | Normal—no apparent changes in breathing, 30–54 breath per minute (BPM), and no cough | 0 |
| | Mild—slightly increased effort breathing, 55–65 BPM, or mild cough | 2 |
| | Moderate—obvious difficulty breathing, 66–80 BPM, or apparent cough | 4 |
| | Severe—respirations labored, open mouth breathing, abdominal breathing, >80 BPM, cyanosis, or hemoptysis | 6 |
| Food consumption | 100% of biscuits | 0 |
| | 10–25% of biscuits remaining | 1 |
| | 25–50% of biscuits remaining | 2 |
| | >50% of biscuits remaining | 3 |
| Fecal consistency | Normal | 0 |
| | Soft | 1 |
| | Fluid | 2 |
| | Fluid and profuse amount | 3 |
| | | **Total** |

**Notes** *(any observed sneezing, vomit, conjunctival erythema, or other abnormalities*)

| **Physical Examination Under Anesthesia** | | |
|---|---|---|
| **Parameter** | **Description** | **Score** |
| Rectal temperature *(taken immediately after sedation)* | Normal (37.0–38.9°C) | 0 |
| | Low-grade fever (39–39.5°C) | 2 |
| | Fever (>39.5°C) | 4 |
| Heart rate | Normal (up to 19 BPM over baseline, ***/***) | 0 |
| | Mild tachycardia (20–39 BPM over baseline), | 1 |
| | Moderate tachycardia (40–69 BPM over baseline) | 2 |
| | Severe tachycardia (>70 BPM over baseline) | 3 |
| Respiratory rate | Normal—30–54 BPM | 0 |
| | Mild tachypnea—55–65 BPM | 2 |
| | Moderate tachypnea—66–80 BPM | 4 |
| | Severe tachypnea—>80 BPM | 6 |
| $SpO_2$ | Normal (95–100%) | 0 |
| | Mildly decreased (90–94%) | 1 |
| | Moderately decreased (87–89) | 2 |
| | Severely decreased (<87) | 3 |
| Body weight | Normal (0–3% loss) | 0 |
| | Mild (4–9% loss) | 1 |
| | Moderate (10–16% loss) | 2 |
| | Severe (>16% loss) | 3 |
| Dehydration | Normal skin turgor, moist mucous membranes | 0 |
| | Skin tenting or dry mucous membranes | 1 |
| | Skin tenting and dry mucous membranes | 2 |
| | Skin tenting, dry mucous membranes, and sunken eyes | 3 |
| | | **Total** |

| |
| --- |
| **Notes** *(auscultation findings if applicable, conjunctival erythema, palpable masses, or any other abnormalities)* |

The sheets above were modified from a previous nonhuman primate influenza A virus study to include clinical signs relevant to COVID-19 and respiratory rates for crab-eating macaques (1-3). Physical examinations were performed whenever an animal was anesthetized.
